# *Ruminococcus torques* is a keystone degrader of intestinal mucin glycoprotein, releasing oligosaccharides used by *Bacteroides thetaiotaomicron*

**DOI:** 10.1101/2024.01.15.575725

**Authors:** Sadie R. Schaus, Gabriel Vasconcelos Periera, Ana S. Luis, Emily Madlambayan, Nicolas Terrapon, Matthew P. Ostrowski, Chunsheng Jin, Gunnar C. Hansson, Eric C. Martens

**Affiliations:** Department of Microbiology and Immunology, University of Michigan, Ann Arbor, MI, USA; Department of Medical Biochemistry and Cell Biology, University of Gothenburg, Gothenburg, Sweden; Centre National de la Recherche Scientifique, Aix-Marseille Univ., UMR7257 AFMB, Marseille, France; Institut National de Recherche pour l’Agriculture, l’Alimentation et l’Environnement, USC1408 AFMB, Marseille, France; Proteomics Core Facility at Sahlgrenska Academy, University of Gothenburg, Gothenburg, Sweden

## Abstract

Symbiotic interactions between humans and our communities of resident gut microbes (microbiota) play many roles in health and disease. Some gut bacteria utilize mucus as a nutrient source and can under certain conditions damage the protective barrier it forms, increasing disease susceptibility. We investigated how *Ruminococcus torques—*a known mucin-degrader that remains poorly studied despite its implication in inflammatory bowel diseases (IBDs)— degrades mucin glycoproteins or their component *O*-linked glycans to understand its effects on the availability of mucin-derived nutrients for other bacteria. We found that *R. torques* utilizes both mucin glycoproteins and released oligosaccharides from gastric and colonic mucins, degrading these substrates with a panoply of mostly constitutively expressed, secreted enzymes. Investigation of mucin oligosaccharide degradation by *R. torques* revealed strong fucosidase, sialidase and β1,4-galactosidase activities. There was a lack of detectable sulfatase and weak β1,3-galactosidase degradation, resulting in accumulation of glycans containing these structures on mucin polypeptides. While the Gram-negative symbiont, *Bacteroides thetaiotaomicron* grows poorly on mucin glycoproteins, we demonstrate a clear ability of *R. torques* to liberate products from mucins, making them accessible to *B. thetaiotaomicron*. This work underscores the diversity of mucin-degrading mechanisms in different bacterial species and the probability that some species are contingent on others for the ability to more fully access mucin-derived nutrients. The ability of *R. torques* to directly degrade a variety of mucin and mucin glycan structures and unlock released glycans for other species suggests that it is a keystone mucin degrader, which may contribute to its association with IBD.

**Importance:** An important facet of maintaining healthy symbiosis between host and intestinal microbes is the mucus layer, the first defense protecting the epithelium from lumenal bacteria. Some gut bacteria degrade different components of intestinal mucins, but detailed mechanisms used by different species are still emerging. It is imperative to understand these mechanisms as they likely dictate interspecies interactions and may illuminate particular species associated with bacterial mucus destruction and subsequent disease susceptibility. *Ruminococcus torques* is positively associated with IBD in multiple studies. We identified mucin glycan-degrading enzymes in *R. torques* and found that it shares mucin degradation products with another gut bacterium implicated in IBD, *Bacteroides thetaiotaomicron*. Our findings underscore the importance of understanding the mucin degradation mechanisms of different gut bacteria and their consequences on interspecies interactions, which may identify keystone bacteria that disproportionately contribute to defects in mucus protection and could therefore be targets to prevent or treat IBD.

## Introduction

The relationship between mammalian hosts and their resident gut microbial communities (microbiota) is both complex and dynamic. To coexist with densely populated and mostly beneficial colonic microbes, appropriate separation needs to be maintained between the microbiota and intestinal tissue. Mammals have evolved a series of physical and immunological defenses to promote this separation. Some bacteria—especially pathogens—can disrupt these defenses and eventually promote disease^1^. Secreted mucus is a critical component of the host defense in the colon and forms a physical barrier between epithelial cells and lumenal bacteria, preventing direct contact. Previous studies have demonstrated the ability of some gut bacteria to affect mucus properties *in vivo,* particularly in the context of a fiber-deficient diet, leading to decreased mucus thickness observed in fixed tissues, increased mucus penetrability and decreased growth rate of the inner mucus layer^2–5^. Increased abundance, activity and/or presence of mucin-degrading bacteria has been associated with detrimental health effects, including increasing pathogen susceptibility^6,7^, development of spontaneous colitis like that which occurs in various inflammatory bowel diseases (IBDs)^8^, increased allergen sensitivity^9^ and higher mortality due to carbapenem-associated graft versus host disease^10^. While the interaction between mucin-degrading bacteria and the status of the mucus barrier is emerging as a facet that influences host health, precise mechanisms of bacterial mucin-degradation are still being determined. Due to the emerging roles of bacterial mucus degradation in multiple diseases, there is a critical need to investigate bacterial mucin-degrading systems to illuminate potential therapeutic targets to prevent this process.

Mammalian colonic mucus is predominantly composed of high molecular weight (∼2.5 MDa) MUC2 monomers, which are disulfide cross-linked to form a polymeric glycoprotein network harboring thousands of structurally diverse, sterically hindered *O*-linked glycans (herein, *O*-glycans) attached to each MUC2 polypeptide^11,12^. The *O*-glycans attached to MUC2 are clustered in two central mucin domains and account for up to 80% of its total mass^13^. The mucin domains are formed by *O*-glycosylation of sequences rich in hydroxy amino acids (serine, threonine) and proline (so called PTS sequences). Though they are composed of only five monosaccharides, there are typically more than 100 unique *O*-glycan structures present, making mucins a highly complex substrate that requires a correspondingly large repertoire of enzymes to degrade^13^. Adding to its complexity, *O*-glycan composition varies both between different segments of the gut and between individuals. In humans, fucosylated glycans decrease in abundance moving from the proximal to distal gastrointestinal tract, while sialylated glycans exhibit an opposite trend^14^. Individual blood group status (A, B, AB, H) is also reflected in secreted *O*-glycans as terminal, non-reducing end sugar linkages^1^.

Alteration in mucin production or *O*-glycan structural diversity has been associated with intestinal disease, particularly IBDs. Conventional mice genetically lacking Muc2 (Muc2^-/-^), a very severe defect not observed in humans, develop spontaneous intestinal inflammation and eventual colorectal cancer, presumably due to direct contact between colonic epithelial cells and gut bacteria^15,16^. Consistent with this, antibiotic treated Muc2^-/-^ mice showed lower levels of proinflammatory cytokines and decreased inflammation compared to those not receiving antibiotic treatment^17^. More subtle changes to MUC2 glycosylation have also been modeled in mice and may reflect polymorphisms that occur in humans. Loss of enzymes for making core 1 and core 3 *O*-glycan base structures, which in turn give rise to many other elongated *O*-glycans present in colonic MUC2, in intestinal epithelial cells of mice leads to spontaneous inflammation in the colon, which is relieved by antibiotic treatment^18^. Polymorphisms in core 1 synthase molecular chaperone (*Cosmc*), an X-linked gene that encodes a chaperone required for C1GalT1 (the β1,3 galactosyltransferase that synthesizes core 1) function, are associated with development of the rare, hematological autoimmune disease Tn syndrome, but are also a risk factor for development of IBDs in males^19^. In mice, *Cosmc* loss in intestinal epithelial cells drives inflammation in males that is dependent on the presence of the microbiota^20^. Individuals experiencing active ulcerative colitis have decreased sulfation and sialylation of their *O*-glycans relative to healthy controls^21^. The association between decreased *O*-glycan length or decreased abundance of terminal “capping” residues and IBD suggests that *O-*glycan complexity is critical to maintaining host health. This also implies that bacteria capable of reducing mucin glycan complexity by degrading these structures may contribute to the development of these diseases.

Historically, porcine gastric mucin (PGM) and PGM-derived substrates have been used to study mucin-degrading abilities of gut bacteria, as they are readily available. Species known to degrade PGM to varying degrees include several species of *Bacteroides* (*B. fragilis, B. vulgatus, B. thetaiotaomicron, B. caccae*)*, Akkermansia muciniphila, Ruminococcus gnavus, Ruminococcus torques, Peptostreptococcus russellii,* and multiple *Clostridiales* and *Bifidobacterium* species^22–24^. While the complex mucin substrates available in the gut provide a potentially rich nutrient source for bacteria, some mucin-degraders can only access certain mucin components. For example, total growth yield of *B. thetaiotaomicron* on PGM glycoprotein is poor^25,26^, but is greatly increased on *O-*glycans that have been chemically released from PGM^27^. Many bacterial carbohydrate active enzymes (CAZymes) active on mucins have been identified^22,28^, although a comprehensive list of bacteria and enzymes involved in mucin degradation is still needed. While less is currently known about bacterial proteases capable of degrading the mucin polypeptide backbone, some have been identified, including M60-like metalloproteases^29,30^.

Despite widespread use of PGM as a substrate to measure bacterial growth on intestinal mucins, studies investigating growth on different, more relevant mucin substrates, especially highly sulfated mucins from the mammalian colon^31^, remain limited^32^. Experimental evaluation of growth on colonic mucins may reveal substantially different and more physiologically relevant enzymatic mechanisms due to their structural differences from gastric mucins. Notably, a previous study found that *R. torques, A. muciniphila, B. bifidum,* and *R. gnavus* all degrade human colonic MUC2, and *R. torques* was the most efficient of the species investigated^33^.

In addition to degrading human MUC2*, R. torques* has also been associated with IBDs in multiple studies. *R. torques* is more prevalent and abundant in people with IBDs, found in 10% of healthy individuals, but 45% of individuals with ulcerative colitis and 38% of individuals with Crohn’s disease, while also being ∼100-fold more abundant in individuals with IBDs^33^. Further, *R. torques* is one of only two bacterial species that is positively correlated with the presence of anti-*Saccharomyces cerevisiae* antibodies, a marker of Crohn’s disease^34^. A recent prospective study that followed first degree relatives of individuals with Crohn’s disease before being diagnosed, found that increased colonization with *R. torques* was the top microbial risk associated with developing Crohn’s disease^35^. While *R. torques* and its metabolism remain generally understudied, previous work demonstrated that *R. torques* ferments PGM and degrades the blood group A and H antigen components of *O*-glycans and Lewis x and y antigens present in intestinal glycosphingolipids^36–38^. Given the implication of *R. torques* as a mucin-degrader associated with IBD^33,35,39^, we aimed to characterize the mechanisms by which *R. torques* degrades both gastric and colonic mucins as well as the role of *R. torques* mucin degradation within a bacterial community.

## Results

### R. torques grows on both mucin glycoproteins and released mucin oligosaccharides

It was previously observed that *R. torques* has the ability to degrade components of porcine gastric mucin (PGM) and human MUC2 using biochemical approaches. However, direct growth measurements of *R. torques* on various mucin and non-mucin substrates have not been reported^33,37^. Since colonic mucin sources are not commercially available, we isolated our own trypsin resistant large (>300 kDa) mucin domain glycopeptides from colonic MUC2 (cMUC2) from pigs^31,40^. PGM and cMUC2 represent the complex form in which mucin glycoproteins are secreted *in vivo,* with *O*-glycans remaining bound to the mucin polypeptide backbone. Using mucin glycoproteins as starting material, we generated chemically released *O-*glycans from each, yielding gastric mucin *O-*glycans (gMO) and colonic mucin *O*-glycans (cMO). These four mucin substrates were used to assess the ability of three different species of gut bacteria, including a human *R. torques* isolate (*R. torques* VIII-239^37^) to degrade different mucin glycoproteins and *O*-glycan components. Examination of gastric and colonic substrates aids us in assessing the ability of bacteria to degrade *O-*glycans of various composition, as there are known variations in abundances of terminal capping residues throughout the gastrointestinal tract^14^.

*Bacteroides thetaiotaomicron* and *Akkermansia muciniphila* are both known to degrade components of gastric mucin^25,27,41^. Interestingly, in our growth experiments they displayed partially opposing preferences for gastric and colonic mucins or their component *O*-glycans, while *R. torques* displayed the broadest mucin-degradation abilities. On the mucin glycoprotein substrates, *A. muciniphila* and *R. torques* grew to a higher max absorbance (600nm) than *B. thetaiotaomicron* (**Figure 1A,B**). In contrast, *R. torques* and *B. thetaiotaomicron* grew to a higher max absorbance on the mucin oligosaccharide substrates than *A. muciniphila* (**Figure 1C,D**). As positive controls for growth, *B. thetaiotaomicron* and *R. torques* were grown on glucose (**Figure 1E**) and *A. muciniphila* was grown on *N*-acetylglucosamine (**Figure 1F**), based on previous growth profiles^6^.

**Figure 1.**
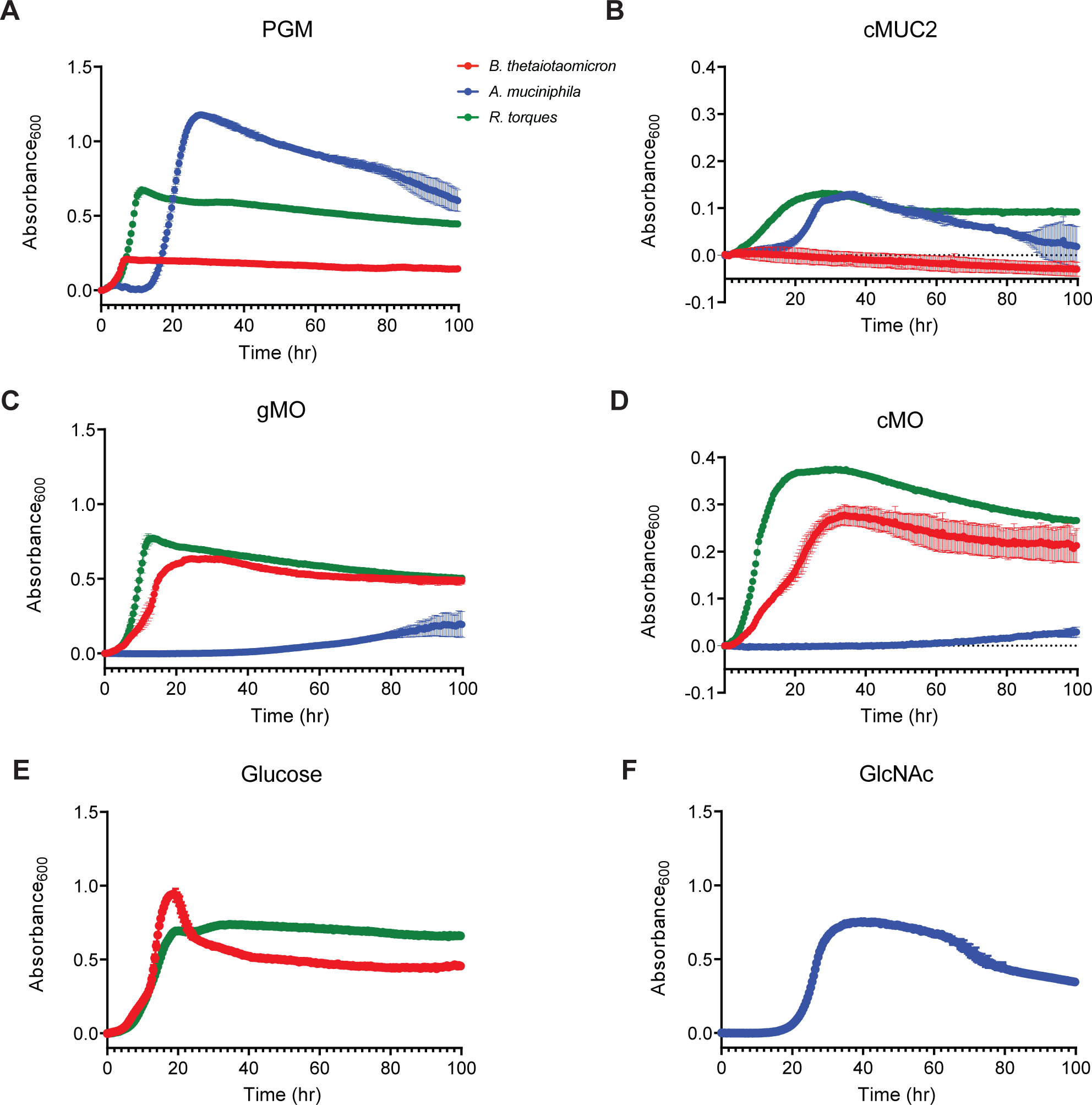
Variation in bacterial growth preferences for different mucin substrates. **A-F**, growth of three commensal gut bacterial species (*Bacteroides thetaiotaomicron, Akkermansia muciniphila,* and *R. torques)* measured anaerobically at 37°C by absorbance at 600 nm on mucin glycoprotein from porcine gastric mucus (**A**, PGM; 10 mg/ml final; n=3), mucin glycoprotein purified from porcine colonic mucosa (**B**, cMUC2, 2.5 mg/ml final; n=3), released oligosaccharides from PGM (**C**, gMO, 10 mg/ml final; n=3), and released oligosaccharides from cMUC2 (**D**, cMO, 5 mg/ml final; n=2). Growths on glucose (**E**, 5 mg/ml final, *B. thetaiotaomicron* and *R. torques,* n=3) and *N*-acetylglucosamine (**F**, GlcNAc, *A. muciniphila,* n=3) were included as positive controls. Each point represents the average of the replicates and error bars represent standard deviation. *B. thetaiotaomicron* was grown in Bacteroides minimal medium, *A. muciniphila* was grown in chopped meat medium, and *R. torques* was grown in YCFA medium (see Materials and Methods for medium formula and references).

In addition to its growth on both gastric and colonic mucin glycoproteins and mucin oligosaccharides, *R. torques* was able to grow on three of the five mucin monosaccharides: galactose, *N*-acetylgalactosamine, and *N*-acetylglucosamine (**Figure S1**). Note that a closely related species, *Ruminococcus gnavus*, has been shown to grow on *N*-acetyl neuraminic acid using a novel *trans-*sialidase mechanism that introduces a 2,7-anyhydro bond, allowing *R. gnavus* to catabolize the cleaved sugar^42^. While we did not directly test for this ability, *R. torques* possesses three genes encoding glycoside hydrolase family 33 (GH33) enzymes, which are candidates for this activity, one of which is discussed more below. This may explain its lack of growth on the unmodified sialic acid (Neu5Ac) tested that lacks the introduced 2,7-anhydro linkage.

Growth experiments using other host- and plant-derived polysaccharides that are expected to transit the gut and reach the colon suggest that *R. torques* is a mucin specialist. Separate growth measurements on a panel of 46 different carbohydrates and monosaccharides^32^ only revealed strong growth (≥0.7 average net A_600_ increase) on glucose, fructose, and the mucin monosaccharides noted above (**Figure S1**). *R. torques* grew modestly on glucosamine and keratan sulfate (∼0.3 and ∼0.15 average net A_600_ increases, respectively) but did not demonstrate growth (≥0.1 net A_600_) on the other substrates tested with the exception of the variable growth on laminarin and pectic galactan from lupin (**Figure S1**). The observation of modest growth on keratan sulfate suggests an inability of *R. torques* to remove sulfate from this substrate, despite growth on colonic mucins, which are also known to be sulfated and share a similar poly-*N*-acetyllactosamine backbone structure^14,43^. Because of its broad ability to degrade mucins and *O*-glycans from different regions of the gastrointestinal tract, we sought to understand the enzyme repertoire that *R. torques* uses to degrade mucins and their component glycans.

### Enzymes in R. torques supernatant degrade gastric and colonic mucin glycoproteins

To further characterize how *R. torques* degrades mucin glycoproteins and to identify factors that influence these activities, we evaluated the ability of whole culture, containing both *R. torques* cells and the associated supernatant, and cell-free supernatant to degrade cMUC2 or PGM. To measure this ability, we employed a PAGE gel assay in which high molecular weight MUC2 domains are visualized by staining with periodic acid-Schiff (PAS) staining (**Figure 2A**). It has previously been observed that *B. thetaiotaomicron* and other mucin-degrading *Bacteroides* exhibit substrate-specific activation of genes involved in utilizing *O*-glycans and other polysaccharides^6,27,44^ Thus, we first sought to understand whether the substrate that *R. torques* was grown on influences its ability to degrade cMUC2. Both whole culture and supernatant from *R. torques* grown on either glucose or gMO degraded cMUC2 (**Figure 2A,B,C; Figure S2A,B**). Notably, the glucose-grown supernatant samples degraded 72.0% of cMUC2 (**Figure 2C**), suggesting that *R. torques* constitutively produces sufficient mucin-degrading enzymes to reduce the molecular weight or glycosylation of the mucins tested, even when the cultures they are derived from were grown in the absence of mucin substrates. Further, this result suggests that these enzymes are secreted or dissociate from the cell frequently enough to observe activity in the supernatant. Surprisingly, prior growth on gMO significantly decreased cMUC2 degradation compared to prior growth on glucose in both culture and supernatant samples (**Figure 2B,C**). This suggests that growth on gMO suppresses the production or activity of critical mucin-degrading enzyme(s) or another associated function. Degradation of cMUC2 by *R. torques* was inhibited by EDTA, suggesting that key mucin-degrading enzymes are dependent on a metal cofactor (**Figure 2D, Figure S2C**).

**Figure 2.**
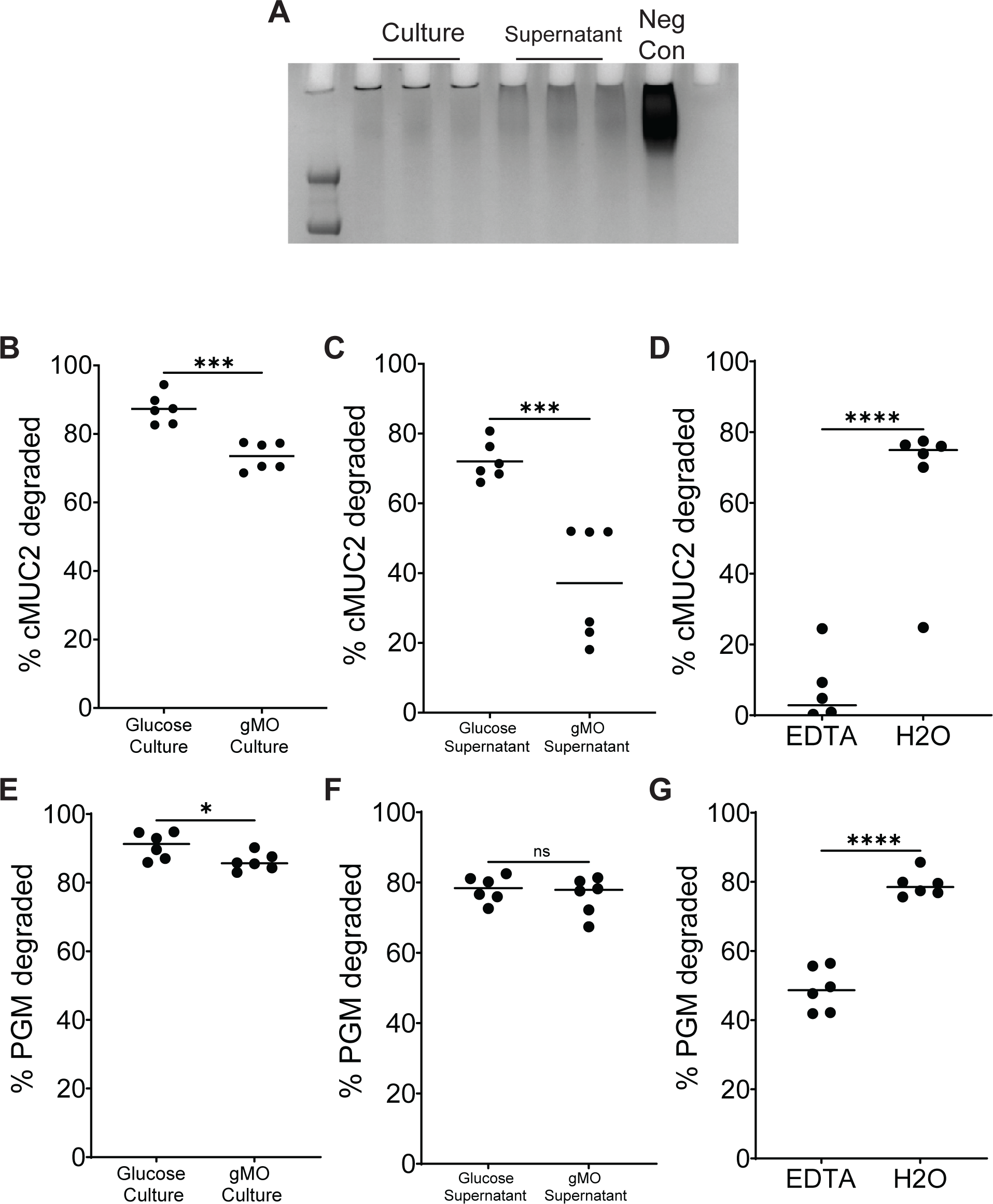
*R. torques* cultures and cell-free supernatants degrade gastric and rectal mucin glycoprotein. **A**, representative periodic acid-Schiff stained gel of samples of *R. torques* culture or supernatant degradation of cMUC2 after growth on glucose. Neg con contains cMUC2 and YCFA medium. **B-C.** Percent colonic mucin glycoprotein (cMUC2) degraded after *Rt* growth in glucose or gastric mucin oligosaccharides (gMO). Culture samples (**B**) or cell-free supernatants (**C**) from these cultures were exposed to cMUC2 for 48 hours. cMUC2 degradation was quantified by visualizing the cMUC2 *Rt* digest on a 4-12% Bis-tris gel, staining with periodic acid-Schiff stain, and quantifying densitometry of the resulting bands using ImageJ. **D**, Percent cMUC2 degraded by *Rt* supernatants in the presence or absence of 10 mM EDTA. **E-F**, Percent gastric mucin glycoprotein (PGM) degraded after by *Rt* culture (**E**) or cell-free supernatant (**F**) after previous growth on glucose or gMO. **G**, Percent PGM degraded by *Rt* supernatants in the presence or absence of 10 mM EDTA. Statistics were analyzed with unpaired, two-tailed t tests. *p<0.05, ***p<0.001, ****p<0.0001, ns=not significant.

PGM glycoprotein was degraded similarly to cMUC2 by both whole culture and cell-free supernatant samples. While prior growth on glucose resulted in culture samples degrading significantly more PGM than gMO-grown cultures, there was no significant reduction in PGM degradation by gMO-grown supernatants compared to glucose-grown supernatants (**Figure 2E,F; Figure S2D**). EDTA also inhibited PGM degradation, although inhibition was not as severe as the effect observed with cMUC2 (48.9% PGM degraded after EDTA treatment, 6.4% cMUC2 degraded after EDTA treatment) (**Figure 2G, Figure S2E**). The variation in effects of EDTA inhibition and prior growth on gMO between cMUC2 and PGM degradation suggest that while there is likely overlap with some shared enzymes that target both substrates, there may be enzymes more critical for cMUC2 degradation that are suppressed after growth on gMO and are more sensitive to loss of metal cofactors.

### Transcriptomic and proteomic approaches to identify secreted R. torques mucin-degrading enzymes

The observation that culture supernatant from *R. torques* cells grown on glucose contain enzymes that degrade cMUC2 suggests that mucin-specific cues are not required for activation of the genes encoding these functions, however prior exposure to gMO can potentially suppress key enzymes. To explore this further, we measured gene expression changes between *R. torques* grown on glucose compared to gMO. *R. torques* was grown to mid-log phase on glucose or gMO, total RNA was isolated and depleted of ribosomal RNA, and transcripts were analyzed by RNA-sequencing. Compared to other mucin-degrading species like *B. thetaiotaomicron*, which alters expression of hundreds of genes in response to growth on *O*-glycans compared to glucose^27^, there were relatively few *R. torques* genes differently regulated on these two substrates. Only 25 genes were upregulated and 35 genes were downregulated during growth in gMO relative to the glucose reference (≥5-fold change; p<0.05) (**Figure 3A, Table S1**).

**Figure 3.**
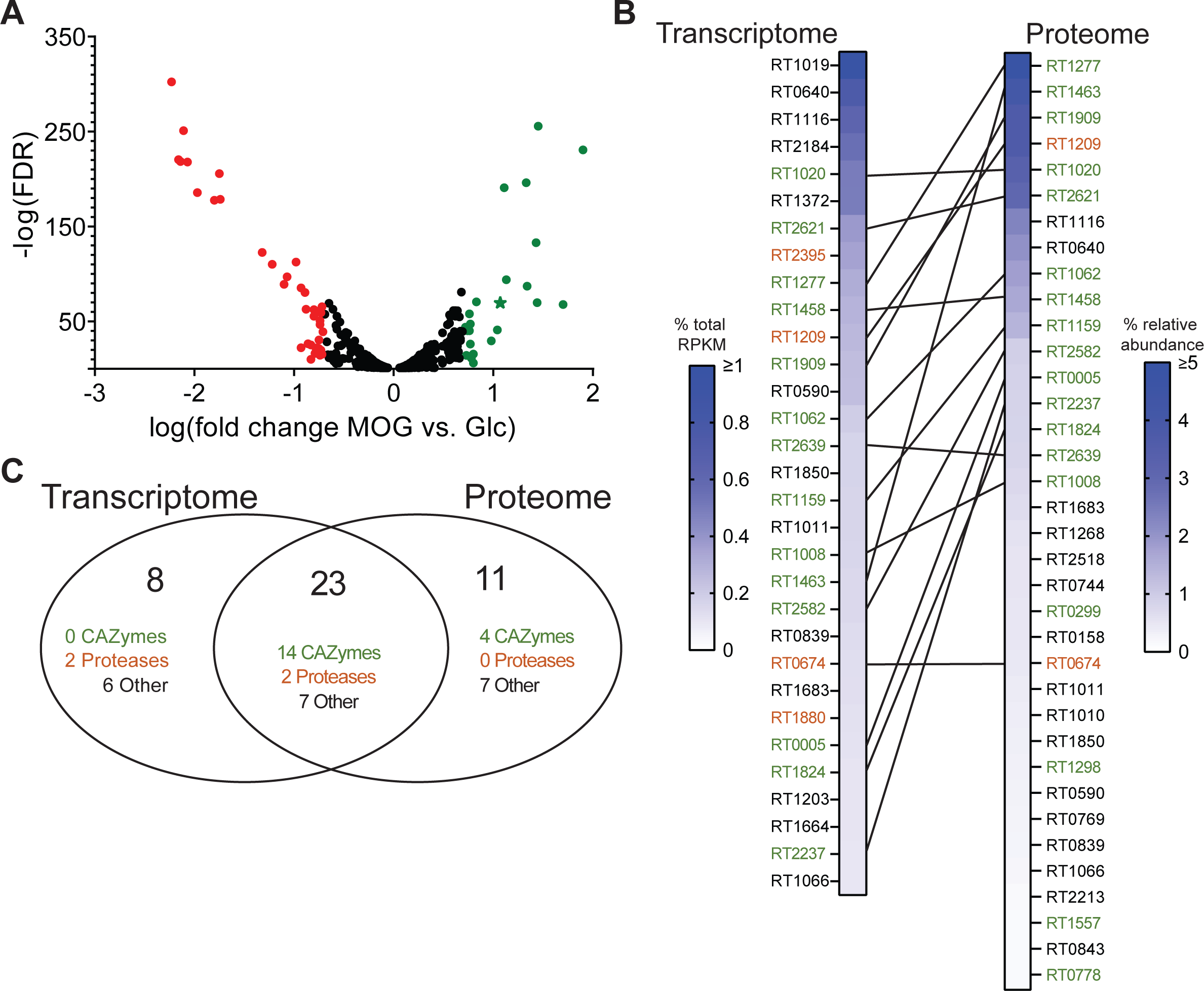
Candidate secreted mucin-degrading enzymes identified in the transcriptome and proteome of *R. torques* despite little transcriptional regulation in response to gastric mucin *O-*glycans. **A**, genes significantly differentially expressed (p<0.05) from RNA-sequencing after growth of *R. torques* on glucose or MOG. Points in green indicate genes upregulated (>5-fold) on MOG versus glucose; points in red indicate genes downregulated (>5-fold) on MOG versus glucose. Green star indicates predicted β-galactosidase. **B**, top highly abundant genes or proteins from the *R. torques* transcriptome (RNA-seq, genes with predicted signal peptide and >0.1% of total RPKM) or proteome (LC-MS/MS, proteins with predicted signal peptide and >0.1% average relative abundance). Lines connecting hits from the transcriptome and proteome indicate genes/proteins present as top hits in both datasets. Green locus tags represent predicted CAZymes or CBM-containing proteins, orange locus tags represent predicted proteases. Transcriptome data n=3; proteome data n=6. **C**, A Venn diagram showing combined top hits from transcriptome and proteome datasets.

Given the observation that supernatants from *R. torques* grown in glucose were capable of degrading cMUC2, it was unsurprising that we detected little difference in expression of genes encoding CAZymes between gMO and glucose grown cells. Only a single putative mucin-degrading CAZyme (rumtor8_01403, predicted GH2 β-galactosidase) was ∼12-fold upregulated during growth in gMO compared to glucose. This putative β-galactosidase was more highly transcribed under gMO conditions and does not contain a predicted signal peptide, thus it was not included in further analysis of *R. torques* secreted mucin-degrading enzymes. Among the observed 35 genes that were upregulated and 25 that were downregulated, most are unlikely to be directly involved in mucin glycan-degradation. These include genes related to ATP metabolism (20% of upregulated genes), bacteriophage (12% upregulated genes, 26% downregulated genes), amine/amino acid metabolism (20% upregulated genes, 14% downregulated genes), nucleotide metabolism (9% downregulated genes), and sugar metabolism (16% upregulated, 3% downregulated) (**Table S1**). Interestingly, some of these genes differentially expressed on mucin *O-*glycans include predicted transporters (12% of upregulated genes, 11% of downregulated genes), including genes predicted to encode sugar transporters(1 upregulated, 1 downregulated) and amino acids (1 downregulated), which could be involved in import of degraded mucin glycoprotein products.

As a complementary approach to identify functions involved in mucin glycan-degradation, we precipitated proteins from *R. torques* supernatants after growth on glucose and analyzed them by LC-MS/MS. Due to an observation that filtered (0.22 μM) *R. torques* supernatants displayed variable defects in degradation of cMUC2 compared to supernatants captured by centrifugation alone (**Figure S3**), samples of both unfiltered, centrifuged supernatants (n=2) and filtered supernatants (n=4, 2 independent experiments) were analyzed. We hypothesized that filtration may result in the removal of critical enzyme(s) necessary for cMUC2 degradation. Samples for each preparation type and experiment were prepared and analyzed in duplicate, and the average relative abundance of each detected protein was calculated for each duplicate pair. We first focused our analysis on proteins containing a predicted signal peptide due to the observed mucin-degrading activity of *R. torques* supernatants (**Figure 2C,F**) and to eliminate contaminants derived from lysed cells. We compiled a list of highly expressed proteins, including all proteins predicted to be secreted that were observed >0.1% relative abundance in at least two of the sample sets (**Table 2**).

Interestingly, there was close similarity between the highly expressed proteins in the filtered samples and the unfiltered samples. There were no putative secreted CAZymes above the 0.1% relative abundance threshold in the unfiltered samples that were not also above this threshold in at least one of the filtered sample groups. This suggests that any observed defect in cMUC2 degradation related to sample filtration may be more likely due to disruption of protein structure, such as large, multi-enzyme complexes, rather than the removal of a particular key enzyme. Thus, we proceeded to include any protein with a predicted signal peptide and representing >0.1% relative abundance in at least 2/3 sample groups as a putative mucin-degrading protein, yielding a list of 34 highly abundant proteins in *R. torques* supernatants, including 18 CAZymes, many of which contain carbohydrate binding modules (CBM) domains, and 2 proteases (**Figure 3B, Table S2**).

To corroborate the proteomics results, we used our transcriptomic data from *R. torques* grown on glucose to identify highly transcribed genes that are also predicted to encode a signal peptide. The top hits representing >0.1% of all transcribed genes based on percent of total RPKM, and are predicted to be secreted, included 31 genes, including 14 CAZymes and 4 proteases (**Figure 3B**). Importantly, most of these top hits were shared with the proteomic data set, including all 14 CAZymes and 2 of the 4 proteases (**Figure 3C, Table S3)**. These enzymes, in addition to the 4 highly expressed CAZymes that are unique to the proteomics dataset, represent top candidates for secreted mucin glycan-degrading enzymes in *R. torques*.

While our results suggest a role for secreted enzymes in *R. torques* mucin-degradation, we assessed all predicted CAZymes in the *R. torques* genome regardless of presence of a signal peptide and their abundance in the transcriptomics and proteomics datasets. We identified two putative CAZymes lacking a predicted signal peptide but present >0.1% relative abundance in the proteomics dataset and/or >0.1% of total RPKM during growth on glucose from the transcriptomics dataset: a carbohydrate esterase family 9 (CE9) and a glycoside hydrolase family 112 (GH112). The CE9 was detected >0.1% of the total relative abundance in all three proteomics sample groups and was just below the >0.1% of total RPKM threshold (0.096%), and the GH112 was above this threshold in two of the three proteomics sample groups and represented 0.11% of the total RPKM. CE9 enzymes are known to deacetylate *N*-acetylglucosamine-6-phosphate, and GH112 enzymes are galacto-*N*-biose and lacto-*N*-biose phosphorylases^45^. As these predicted substrates or products are related to glycan structures in mucins, it is possible that these enzymes are involved in mucin metabolism by *R. torques* and may contribute to the previously observed phenotype that culture samples degraded a larger percentage of cMUC2 than supernatant samples (**Figure 2A, B**).

### R. torques culture supernatants contain large, multidomain enzymes that cleave O-glycan linkages

Analysis of the 18 most highly expressed CAZymes from the transcriptomic and proteomics data sets revealed putative catalytic domains belonging to 12 different CAZy families (**Figure S4**). Predicted activities based on previously characterized enzymes from each of these families reflect expected catalytic roles in degrading mucin *O*-glycans: α− and β− *N*-acetylglucosaminidases (GH89 and GH84, GH20), endo- and exo-α-*N*-acetylgalactosaminidases (GH101 and GH31_18, GH36), sialidase (GH33), endo-β-*N*-acetylglucosaminidases (GH73), β-galactosidases (GH2), and α−fucosidases (GH29, GH95)^32^. Notably, 12/18 of these enzymes are predicted to contain F5/8 type C domains, which are likely carbohydrate binding modules related to the CBM32 family^46^. We often observed multiple repeats of these putative CBMs in a single polypeptide, which may play a critical role in mucin *O-*glycan binding^33^. Many of these enzymes are also notably large, with 17/18 being >125 kDa, and 8/18 >200 kDa (**Table S3**). Despite strong *R. torques* growth on and degradation of sulfated cMUC2 and cMO, as well as previous findings that sulfatases were indispensable for utilization of colonic mucus in *Bacteroides thetaiotaomicron*^31^, there was a surprising lack of predicted sulfatase enzymes. Indeed, bioinformatic homology searches against the SulfAtlas HMM library failed to identify any candidate sulfatases in the *R. torques* genome.

Because of the large size of enzymes detected in *R. torques* supernatants and poorly successful attempts to express and purify recombinant forms of individual enzymes (not shown), we empirically assessed the collective catabolic abilities of the enzymes present in the secretome. Proteins present in *R. torques* supernatants after growth on glucose were precipitated using ammonium sulfate and this mixture was incubated with defined sulfated monosaccharides or small oligosaccharides, many of which are known components of longer mucin *O*-glycans. In accordance with a lack of predicted sulfatase enzymes, we did not observe activities of *R. torques* precipitated proteins on sulfated monosaccharides (**Figure S5**). We did observe sulfatase activity in positive control sonicated *B. thetaiotaomicron* cultures after exposure to keratan sulfate to induce these functions, which included cleavage of GalNAc-6S and GlcNAc-6S that were previously reported for this species^31^ (**Figure S5**). Cleavage by the *R. torques* precipitated proteins was observed with the disaccharide *N*-acetyllactosamine (LacNAc), which presents a terminal, non-reducing end β1,4-linked galactose (**Figure 4A, Figure S6A**). Interestingly, cleavage of lacto-*N*-biose (LNB), a disaccharide isomer of LacNAc with a β1,3 galactose linkage was weaker. Due to the presence of a species with a similar retention time to galactose in control reaction mixtures with LNB (**Figure S6B,H**), the concentration of galactose detected in the LNB negative control by HPAEC-PAD was subtracted from galactose concentrations calculated for LNB samples treated with *R. torques* enzymes. From the concentrations of galactose and *N*-acetylglucosamine released, weaker activity was calculated for LNB, suggesting that *R. torques* is less proficient at removing terminal galactose when present as a β1,3 linkage (**Figure 4A, Figure S6A,B,H**).

**Figure 4.**
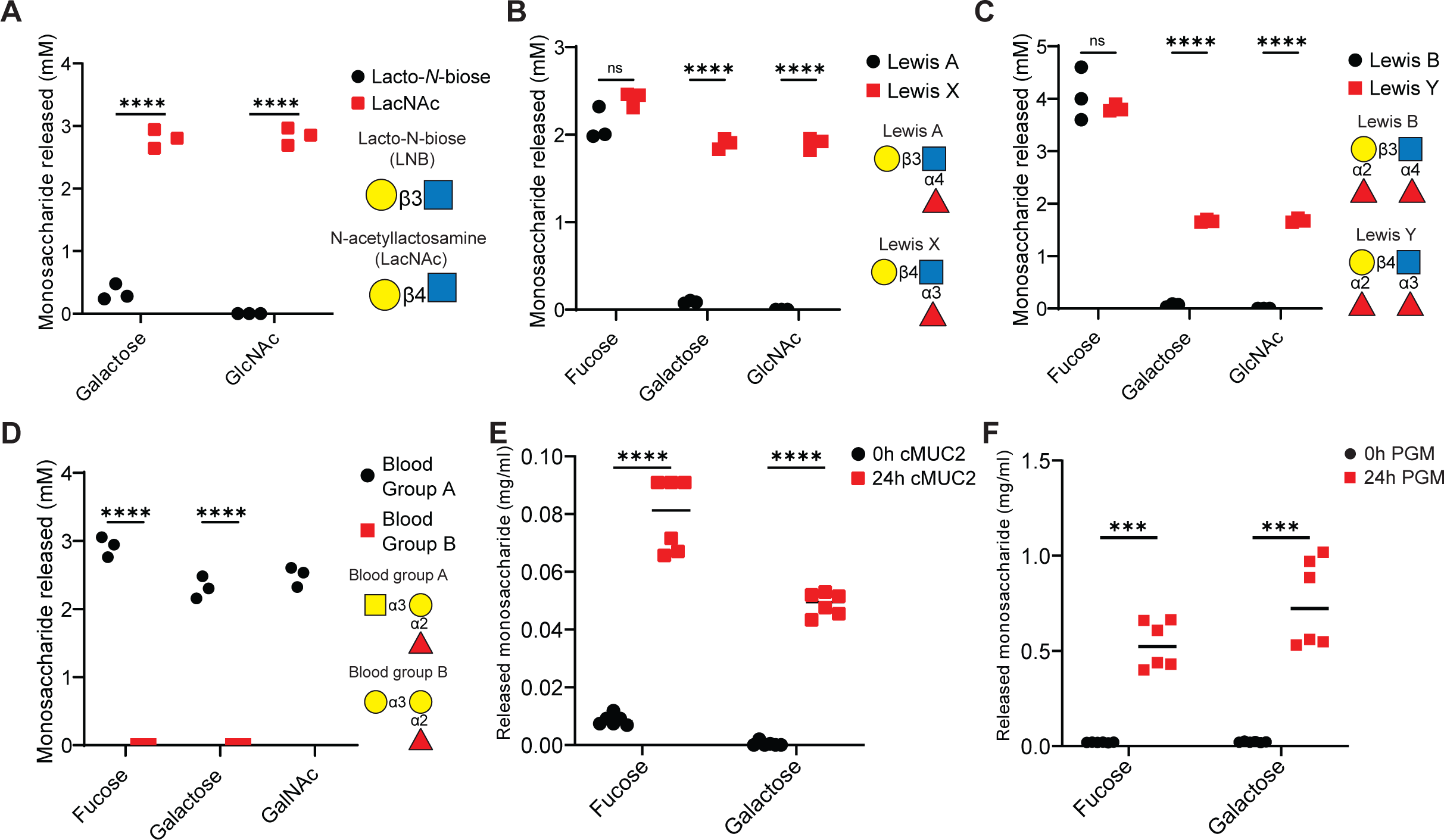
*R. torques* supernatant enzymes exhibit linkage specificity on model mucin glycans and release monosaccharides from intact mucin glycoprotein substrates. **A-D**, quantification of monosaccharides released from Lacto-*N*-biose (LNB) or *N*-acetyllactosamine (LacNAc) (5 mM concentration each, **A**), Lewis A or Lewis X antigens (2mM final concentration each, **B**), Lewis B or Lewis Y antigens (2 mM final concentration each, **C**), and blood group A or blood group B (2 mM final concentration each, **D**) by ammonium sulfate precipitated proteins from *R. torques* supernatants after growth on glucose, measured by High Performance Anion Exchange Chromatography with Pulsed Amperometric Detection. Concentration values were calculated from a standard curve, and calculations resulting in a negative concentration are displayed as zero and were reflected as zero in statistical analyses. For lacto-*N*-biose (LNB) samples, the galactose concentration found in the negative control was subtracted from concentrations calculated for supernatant reactions due to the presence of a species with a similar retention time in the no enzyme control reaction. Statistical analyses were performed with unpaired, two-tailed t tests comparing release of each monosaccharide between substrates in each panel. ns=not significant, ***p<0.001, ****p<0.0001. **E-F**, concentration of fucose or galactose released after 24-hour incubation of *R. torques* supernatants with cMUC2 (2.5 mg/ml final; **E**) or PGM (10 mg/ml final; **F**). Calculated values resulting in a negative concentration are displayed as zero and were recorded as zero for statistical analyses. Statistics were analyzed with paired, two-tailed t tests comparing release of each monosaccharide between substrates in each panel. ***p<0.001, ****p<0.0001.

Fucose was released efficiently from all four fucosylated substrates tested, including Lewis a, b, x, and y (**Figure 4B,C; Figure S6C-F**). Thus, *R. torques* enzymes have fucosidase activities on all mucin glycan-relevant fucose linkages (fucose–α1,2-galactose, fucose–α1,3-*N*-acetylglucosamine, fucose–α1,4-*N*-acetylglucosamine), with no apparent linkage preference as there was no significant difference between the concentration of fucose released when comparing across substrates (**Figure 4B,C**). The trend of enhanced activity on galactose-β1,4-*N*-acetylglucosamine (type 2 chain) versus galactose-β1,3-*N*-acetylglucosamine (type 1 chain) is also apparent with these substrates, as more galactose was released from Lewis x and Lewis y compared to Lewis a or Lewis b (**Figure 4B,C**). We tested activity on two additional terminal *O-* glycan epitopes, blood groups A and B^43^. Strong activity was observed on blood group A, but not blood group B (**Figure 4D, Figure S7A-D**).

While utilizing simplified model oligosaccharides as substrates for *R. torques* supernatants allowed us to probe the presence of activities that cleave specific linkages, we sought to understand whether monosaccharides were also released from the more complex mucin glycoproteins. *R torques* supernatants collected after growth on glucose were incubated with cMUC2 or PGM and samples were collected immediately (0h) and after 24h of anaerobic incubation. Released fucose and galactose were quantified using ultraviolet spectrophotometric assays (Megazyme). The concentrations of both fucose and galactose were significantly increased after 24h incubation with *R. torques* supernatants compared to the 0h samples in cMUC2 and PGM (**Figure 4E,F**). When the concentration of each monosaccharide released after 24 hours was normalized to the respective substrate concentration in each reaction (2.5 mg/ml cMUC2 or 10 mg/ml PGM), less fucose and galactose were released from cMUC2 (fucose: 0.032 ± 0.005; galactose: 0.020 ± 0.002) than from PGM (fucose: 0.053 ± 0.012, p=0.0027; galactose: 0.075 ± 0.023, p=0.0001). This may be due to increased sulfation of cMUC2 glycans relative to PGM glycans^14^, which may inhibit access of *R. torques* due to its observed lack of sulfatase activities (**Figure S5**).

Interestingly, fucose and galactose were also released by *R. torques* supernatants previously grown on gMO from cMUC2 and PGM (**Figure S8A,B**). This suggests that the previously observed decrease in mucin degradation (**Figure 2B,C,E,F**) from gMO-grown supernatants are not due to a loss of the ability to release these two monosaccharides from cMUC2 or PGM. Thus, we have demonstrated the presence of fucosidase and galactosidase activities in the *R. torques* secretome, which are active on both free oligosaccharides and mucin glycoprotein domains.

### Digestion of mucin glycans during R. torques growth

To extend our knowledge of *R. torques* enzymes that cleave linkages present in mucin glycans, we sought to identify specific structures within the mucin glycoprotein substrates that are degraded during growth. We were also interested in identifying structures that were recalcitrant to degradation, suggesting the absence of certain activities, which could also become byproducts available for other members of the microbiota. *R. torques* was grown on PGM or cMUC2 and samples were captured either immediately after inoculation (pre-growth, control) or after growth into stationary phase (post-growth) for each substrate. Residual glycoprotein samples were immobilized on a PVDF membrane to remove any free glycans or media components. The retained glycans that remained attached to the polypeptide backbone after degradation were subsequently released by reductive β-elimination and analyzed using LC-MS/MS. Because of the extreme difficulty in separating free glycans from cultivated medium (data not shown), gMO and cMO were not included in this study.

First, we examined each of the glycans present pre- and/or post-growth, excluding peeling reaction products generated after the reductive β-elimination step, and measured the prevalence of three structural features: sulfation, sialylation, and fucosylation (**Figure 5A**). As expected from known structural variations in mucus along the length of the gut^14^, cMUC2 contained a large proportion of sulfated and sialylated glycans (**Figure 5A**). We measured the mass-to-charge ratio (m/z), deduced structure (when possible) and relative abundance of the detected glycans to determine whether individual glycans were degraded or accumulated. For this analysis, putative peeling reaction products (identified in **Figure S9**) were included as they are derived from intact glycans originally present in the sample and still provide information about the presence of these structural features despite having been further degraded at the reducing end (opposite the typical sites of sulfation, sialylation and fucosylation) in sample processing steps.

**Figure 5.**
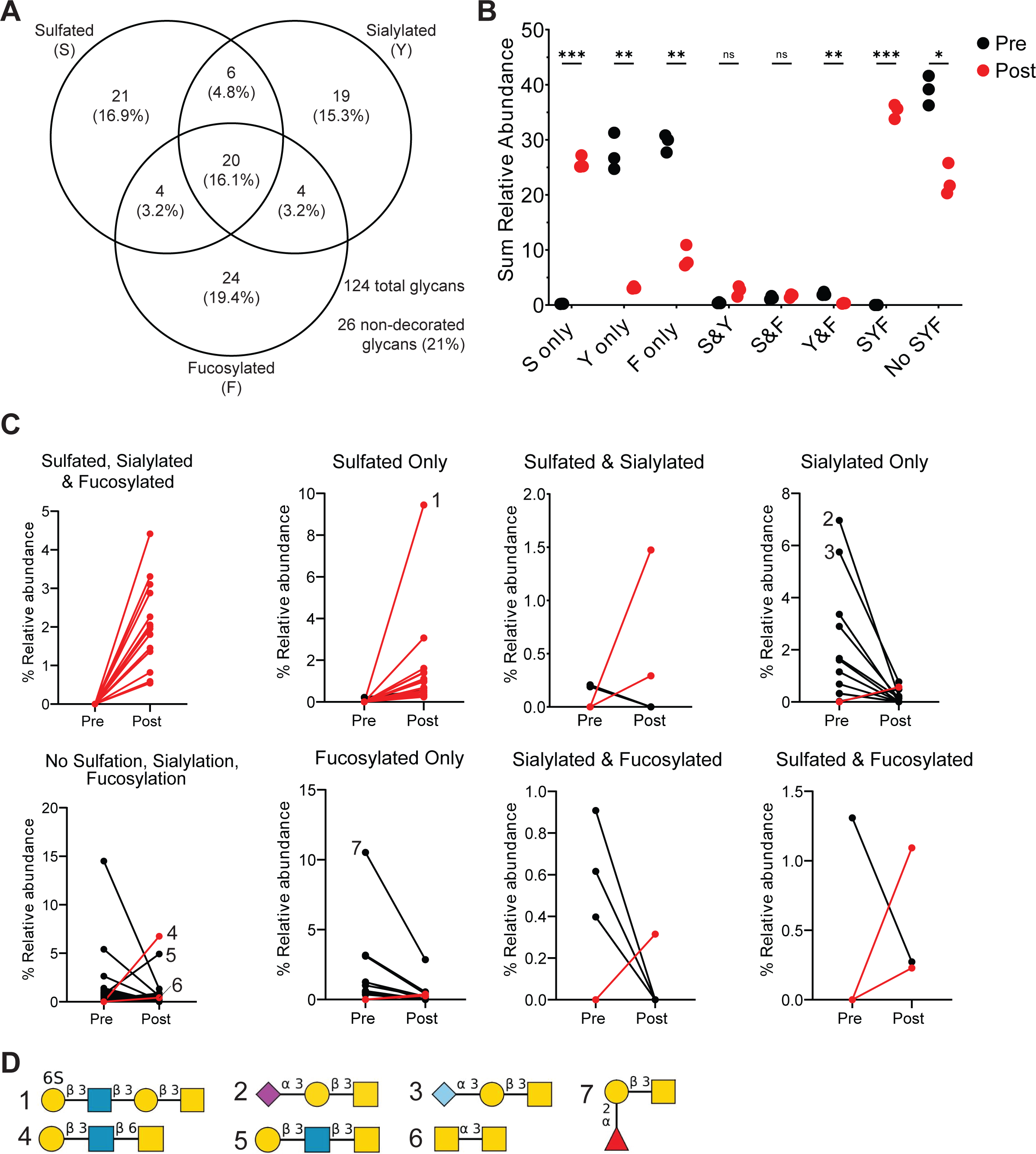
Fucosylated and sialylated glycans are degraded after growth of *R. torques* on cMUC2, but sulfated glycans accumulate. **A**, Number and percentage of total glycans detected in cMUC2 pre- and/or post-growth of *R. torques* displaying indicated structural features by LC-MS/MS. **B**, Sum relative abundance of glycans and inferred peeling reaction products pre- or post-*R. torques* growth on cMUC2 with respective structural features. Statistical analysis was performed with paired, two-tailed t tests between pre- and post-growth samples for each structural feature category. S=sulfated, Y=sialylated, F=fucosylated. **p<0.01, ***p<0.001, ****p<0.0001, ns=not significant. **C**, Relative abundance of glycans detected that were retained on the mucin polypeptide backbone and were significantly different in pre- or post-growth samples of *R. torques* on cMUC2. Red lines indicate glycans that were not detected in any of the pre-growth samples but were detected in the post-growth samples. Each point represents the average relative abundance of three samples. **D**, Putative structures of select detected glycans. Numbers refer to corresponding plot in C. Structures generated with DrawGlycan^63^.

The sum relative abundance of most glycans containing sulfate (*i.e*., those that were sulfated or sulfated, sialylated and fucosylated) increased significantly after *R. torques* degradation of the cMUC2 (**Figure 5B**). The exception to this was glycans that were both sulfated and sialylated or sulfated and fucosylated which showed no change in relative abundance after growth, though these represent small minorities of the total abundance of glycans detected (**Figure 5B**). The accumulation of sulfated glycans further suggests a lack of sulfatase activity in *R. torques.* In contrast, glycans that were sialylated only, fucosylated only, both sialylated and fucosylated, and glycans lacking any of these structural features decreased in sum relative abundance after *R. torques* growth (**Figure 5B**). This suggests strong sialidase and fucosidase activities, the latter of which is concordant with our activity data on model oligosaccharides and mucin glycoprotein substrates (**Figure 4B,C,D,E,F**).

Examining glycans with significantly different relative abundances pre- and post-growth, the majority of glycans that were sulfated, sulfated and fucosylated, or sulfated, sialylated and fucosylated, were undetected before growth and increased in abundance after growth (**Figure 5C, Figure S9A,B,G**). Notably, many of the newly detected sulfated glycans after *R. torques* growth were large (>1 kDa), making it likely that these glycans were present in the initial sample as components of even larger glycans that exceeded the detectable mass range and were partially degraded by *R. torques* to a detectable size (**Table S4**). The accumulation of sulfated glycans was quite remarkable, with one sulfated glycan which was undetectable before growth increasing to >9% average relative abundance after growth (**Figure 5C,D**, structure #1). Indeed, the accumulation of sulfated glycans after *R. torques* growth suggests an inability of *R. torques* to fully degrade sulfated oligosaccharides, leading to their retention on the mucin polypeptide backbone.

In contrast to sulfated glycans, sialylated and fucosylated glycans were more readily degraded by *R. torques*. The majority of glycans significantly different in relative abundance pre- and post-growth that were sialylated, sialylated and fucosylated, or fucosylated only were present in pre-growth samples but decreased in abundance after growth (**Figure 5C, Figure S9F,H,D**). Putative structures from the most highly abundant sialylated glycans pre-growth that decrease in abundance post-growth suggest an ability of *R. torques* to remove both *N*-acetylneuraminic acid (NeuAc) and *N*-glycolylneuraminic acid (NeuGc) from select glycans (**Figure 5C,D**, structures #2 and #3). The deduced structure of a highly abundant fucosylated glycan pre-growth that decreases in abundance post-growth suggests fucose-α1,2-galactose activity (**Figure 5C,D**, structure #7), an activity which we previously observed on Lewis b and y (**Figure 4C**).

Glycans lacking all three structural features (fucosylation, sialylation, sulfation) overall decreased in abundance (**Figure 5B,C; Figure S9C**). Of note, the deduced structures of the two most abundant glycans in this category post-growth present terminal galactose-β1,3-*N*-acetylglucosamine linkages (**Figure 5C,D**, structures #4 and #5). The accumulation of glycans with this terminal linkage further supports a deficiency in enzymes that efficiently cleave this linkage, consistent with the weak activities noted above on Lewis a, Lewis b, and LNB (**Figure 4A,B,C**). There was also an accumulation of *N*-acetylgalactosamine-α1,3-*N*-acetylgalactosamine, suggesting a lack of activity targeting this linkage (**Figure 5C,D**, structure #6). This finding is not surprising, as this putative structure is core 5, a less commonly reported mucin core structure found in human meconium and intestinal cancer^43^.

It is also of note that 16 *N-*glycans were detected pre- and/or post-growth and are included in the above analyses (**Table S4**). Of these, only half were significantly different in abundance between pre-growth and post-growth samples. Overall, *N*-glycans decreased in abundance after growth, representing ∼12% relative abundance pre-growth and ∼6% relative abundance post-growth.

The relative levels of sulfation, sialylation, and fucosylation were also as expected for PGM (*i.e*., fucosylation being much more prevalent), based on known gradients of these structural features throughout the porcine GI tract^14^ (**Figure S10A**). Examining sum relative abundances of glycans and putative peeling reaction products with specific structural features pre- and post-growth indicated a significant decrease in glycans that were sialylated only, fucosylated only, and both sialylated and fucosylated (**Figure S10B**). In contrast, sulfated only and glycans lacking all three structural features increased in abundance post-growth (**Figure S10B**).

One glycan represented a large relative abundance among the sulfated only glycans. This glycan was originally undetected but after growth represented >4% relative abundance of all glycans (**Figure S10C,D,** structure #1**; Table S5, Figure S11D**). The deduced structure of this glycan is predicted to contain an internal *N*-acetylglucosamine-6S linkage. Interestingly, one deduced structure lacking all three features decreased dramatically in abundance post growth, suggesting an ability to release *N*-acetylglucosamine-α1,4-galactose from mucin glycans (**Figure S10C,D,** structure#2; **Figure S11B**). In contrast, the glycan with the predicted structure galactose-β1,3-*N-*acetylgalactosamine (core 1) increased dramatically (from <1% to >20%) in abundance after growth, suggesting deficient activity against this structure (**Figure S10C,D,** structure #3; **Figure S11B**). As observed with cMUC2, the majority of sialylated glycans were degraded, including one glycan predicted to contain a branched core 1-derivative with terminal *N*-acetylneuraminic acid-α2,6-*N*-acetylgalactosamine linkage representing >1.5% relative abundance pre-growth and becoming undetectable after growth (**Figure S10C,D,** structure #4; **Figure S11F**).

While most fucosylated glycans decreased in abundance post-growth, consistent with our previous observations of strong fucosidase activities in *R. torques,* a glycan with a predicted structure of fucose-α1,2-galactose-β1,3-*N*-acetylgalactosamine increased in abundance post-growth (**Figure S10C,D**, structure #5; **Figure S11A**). This is the opposite of the trend observed with the same deduced glycan structure in cMUC2 (**Figure 5C,D**), which may suggest that key enzymes involved in degrading this substrate are not expressed or active during growth on PGM specifically. Alternatively, variation in glycan location and distribution throughout the mucin polypeptide backbone in PGM versus cMUC2 may result in varied steric hindrance and alter access of enzymes to these glycans, which we are unable to assess using this approach. Additionally, 12 *N-*glycans were identified in pre- and/or post-growth samples that were included in our PGM analysis, 8 of which were significantly different in abundance between pre- and post-growth samples. *N-*glycans increased in abundance overall from ∼1.4% relative abundance pre-growth to ∼6.8% relative abundance post-growth.

Collectively, our data indicate that *R*. *torques* efficiently degrades many fucosylated and sialylated mucin glycans, while largely failing to degrade sulfated ones or sulfated regions of larger glycans. These observations from putative structures of glycans retained on the mucin polypeptide backbone after growth, paired with previous observations of supernatant activities on model glycans, provide insight into potential products released for cross-feeding with other members of the microbial community. These results suggest *R. torques* releases monosaccharides (fucose, galactose, *N*-acetylglucosamine, and *N*-acetylgalactosamine) from mucin glycans, while heavily sulfated glycans are retained on the mucin polypeptide backbone.

### R. torques enhances growth of B. thetaiotaomicron on porcine gastric mucin

The results described above provide evidence that *R. torques* liberates monosaccharides and possibly *O*-glycan products from mucin glycans, as well as leaving partially digested *O*-glycans bound to the mucin polypeptide backbone. We next sought to examine whether these products become accessible to other mucin-degraders. *B. thetaiotaomicron* degrades *O-*glycans released from both PGM and cMUC2, but grows to a low maximum absorbance or not at all on cMUC2 and PGM glycoproteins (**Figure 1A,B**)^6,27^. Given the latter observation, we sought to understand interspecies interactions between *R. torques* and *B. thetaiotaomicron* with respect to utilization of PGM (due to limited availability of cMUC2 we chose PGM as the more readily available substrate). Fortuitously, *R. torques* also grows robustly (max A_600_ 0.87±0.01 in 48 hrs of growth) in a partially defined medium (DM) when PGM is present as the major carbohydrate source (**Figure S12**), and this medium is also suitable for growth of *B. thetaiotaomicron* and other *Bacteroides*^48,49^. As *B. thetaiotaomicron* grows comparatively poorly (max A_600_ 0.34±0.01 in 48 hrs) by itself in DM-PGM medium (**Figure S12**), we hypothesized that co-culture with *R. torques* might support enhanced *B. thetaiotaomicron* growth. It is of note, that while the total growth yield of *R. torques* is higher than that of *B. thetaiotaomicron* on PGM*, B. thetaiotaomicron* reaches its maximum growth faster (∼9h) than *R. torques* (∼19h), which may confer a competitive advantage in the ability of *B. thetaiotaomicron* to access shared products when in co-culture with *R. torques.* Despite its lower growth yield on PGM compared to *R. torques*, however, the exponential doubling time of *B. thetaiotaomicron* (1.07 hrs ± 0.04 hrs) on PGM was not significantly different than that of *R. torques* (1.07 hrs ± 0.15 hrs), suggesting that *B. thetaiotaomicron* is able to access less of this substrate, but efficiently utilizes the components it can access.

To test if *B. thetaiotaomicron* profits from the presence of *R. torques* during growth on PGM, both species were cultured together in DM containing either glucose or PGM and were passaged daily for 5 days with a 1:20 dilution factor. Measured by relative abundance, the *B. thetaiotaomicron tdk*^-/-^ strain (a thymidine kinase-deficient parent strain in which allelic exchange mutants are generated^50^) dominated in co-culture using DM-glucose medium, quickly becoming >91% of the community at each timepoint (**Figure 6A**). Surprisingly, in daily DM-PGM co-cultures, *B. thetaiotaomicron tdk*^-/-^ also dominated the community, representing >76% relative abundance at each timepoint, and this dominance remained stable through day 5 (**Figure 6B**). The dominance of *B. thetaiotaomicron* in co-culture with *R. torques* despite its comparatively poor ability to grow in monoculture on PGM suggests that *R. torques* liberates products from PGM that *B. thetaiotaomicron* can access.

**Figure 6.**
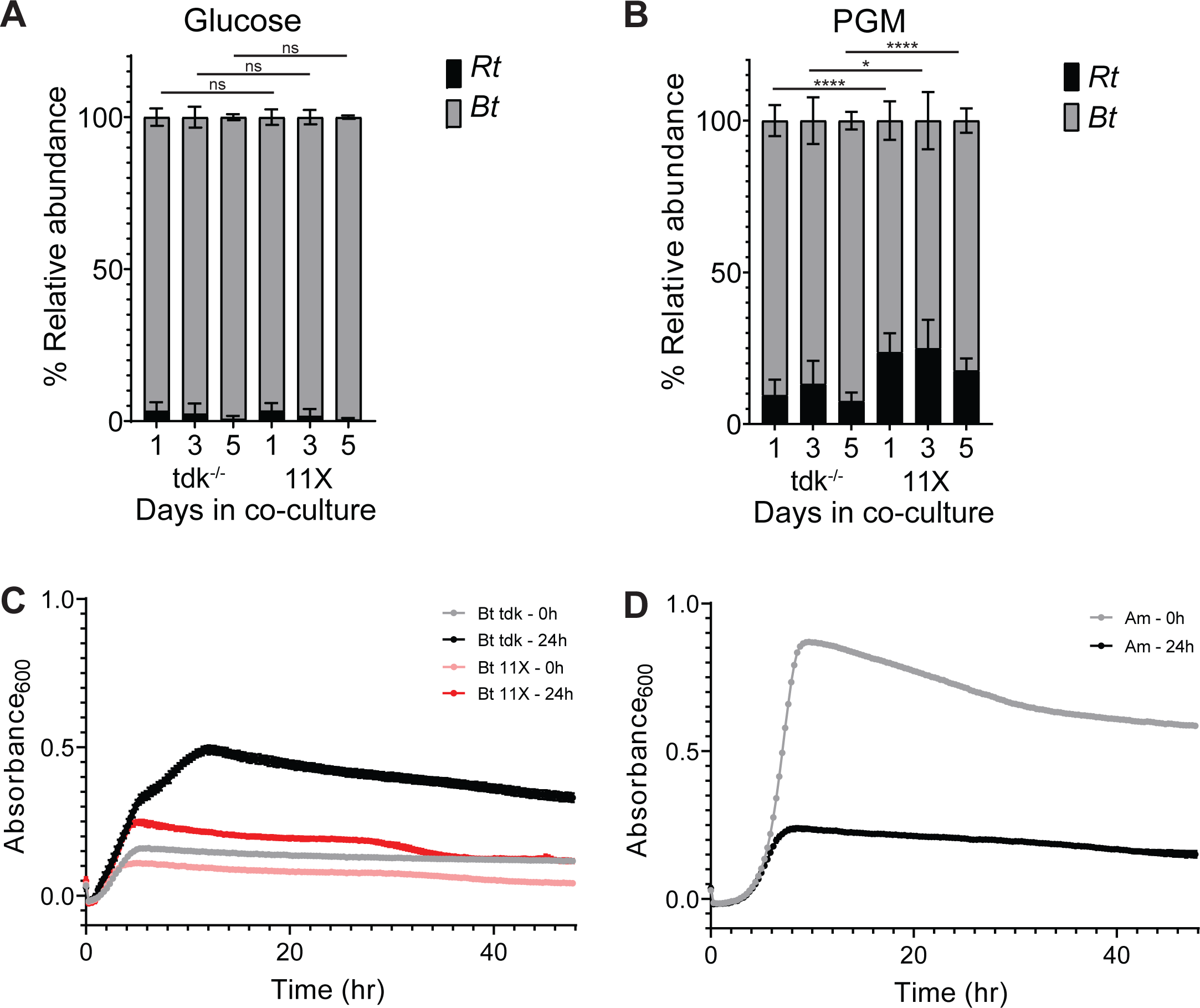
*R. torques* generates monosaccharide and *O-*glycans products from PGM that *Bacteroides thetaiotaomicron* can utilize. **A-B**, relative abundance of *R. torques* and *B. thetaiotaomicron tdk*^-/-^ or 11X mutant in co-culture grown in partially defined medium (DM) and glucose (**A**, 5 mg/ml) or PGM (**B**, 10 mg/ml) calculated by qPCR. Statistics were analyzed with two-tailed, unpaired t tests comparing relative abundance of *R. torques* vs. *B. thetaiotaomicron tdk*^-/-^ co-culture to relative abundances in *R. torques* vs. *B. thetaiotaomicron* 11X mutant co-culture at each respective timepoint. *=p<0.05, ****=p<0.0001, ns=not significant, n=9. **C-D**, growth of *B. thetaiotaomicron* (*tdk*^-/-^, 11X mutant, **C**) and *A. muciniphila* (**D**) on PGM pre-digested by *R. torques* supernatant for 0h or 24h. *B. thetaiotaomicron* was grown in *Bacteroides* minimal medium and *A. muciniphila* was grown in chopped meat medium. N=3, points represent mean ± standard deviation.

An additional benefit to exploring the interactions between *R. torques* and *B. thetaiotaomicron* is that the latter is genetically tractable. Many genes in this species have previously been associated with *O*-glycan degradation^27,31,51^ and in some cases deleted to produce mutants that are deficient at using *O*-glycans *in vitro* or *in vivo*^10,31^. We next sought to assess whether the consumption of released mucin *O-*glycans—as opposed to consumption of liberated monosaccharides like fucose and galactose—is a significant aspect of the ability of *B. thetaiotaomicron* to persist in co-culture with *R. torques* in medium containing PGM. To test this, we co-cultured *R. torques* and a *B. thetaiotaomicron* mutant, which lacks 93 genes spanning 11 PULs (*B. thetaiotaomicron* 11X mutant)^10^. Importantly, this mutant has a large growth defect on gMOs and *N*-acetylgalactosamine, but not on *N*-acetylglucosamine, galactose, fucose, or glucose (**Figure S13A-F**). Compared to co-cultures with the parent *B. thetaiotaomicron tdk*^-/-^, the mucus-deficient 11X mutant strain has no defect in co-culture with *R. torques* on glucose but shows a significant defect on PGM (**Figure 6A,B**). This suggests that *B. thetaiotaomicron* relies on its ability to utilize *O-*glycans and/or free *N*-acetylgalactosamine through the 11 missing systems to access some of the PGM products released by *R. torques*.

To further demonstrate that *B. thetaiotaomicron* persistence during co-culture with *R. torques* on PGM is due to liberation of previously inaccessible glycans, we isolated the enzyme-containing supernatant after *R. torques* was grown on glucose and incubated it with PGM. A sample was collected and frozen immediately, to control for the presence of other metabolites in the spent media supernatant. Another sample was allowed to digest the PGM for 24 hours. After boiling, centrifugation, and dialysis of both substrates (1 kDa cutoff, to remove monosaccharides, **Figure S13G,H**), the digested PGM was sterilized and used as a growth substrate for *B. thetaiotaomicron.* In medium supplemented with the 0-hour control digest, *B. thetaiotaomicron* growth (Max A_600_ in 48 hours 0.160 ± 0.004) was similar to untreated PGM (Max A_600_ in 48 hours 0.212 ± 0.008) (**Figure 1A,6C**). Total growth of *B. thetaiotaomicron* increased dramatically on the *R. torques* supernatant digested PGM (**Fig. 6C**). This further suggests that *R. torques* releases products from PGM that *B. thetaiotaomicron* can use for growth but cannot access without the help of *R. torques*. Since the PGM digest substrate was dialyzed to remove monosaccharides, *R. torques* must be releasing some *O-*glycan fragments during growth in addition to free monosaccharides. While the growth of the mucus-deficient *B. thetaiotaomicron* 11X mutant increased on the 24-hour *R. torques* PGM digest relative to the 0-hour control digest, the average maximum absorbance values of the *B. thetaiotaomicron* 11X mutant after growth on the 24-hour digest (0.247 ± 0.008) were only 50.3% of the average maximum absorbances of *B. thetaiotaomicron tdk*^-/-^ (0.492 ± 0.013) (**Figure 6C**). This further suggests that the products released by *R. torques* from PGM include *O-*glycans, as the 11X mutant has a strong growth defect on gMO (**Figure S13A**).

While our results support a model of cross-feeding between *R. torques* and *B. thetaiotaomicron* on PGM in part through oligosaccharides released by *R. torques*, they suggest a different relationship exists between *R. torques* and *A. muciniphila.* Similar to *R. torques*, *A. muciniphila* grows to a high maximum absorbance value on undigested PGM (**Figure 1**). However, *A. muciniphila* displayed decreased total growth on *R. torques* digested PGM compared to undigested PGM (**Figure 6D**), suggesting that while *R. torques* and *A. muciniphila* both are mucin glycoprotein-degraders, *R. torques* products released from PGM are an insufficient growth substrate for *A. muciniphila*. This suggests that each species uses different mechanisms to release unique products and that the products released from PGM by *R. torques* are not an optimal nutrient source for *A. muciniphila*, which is consistent with its weaker growth phenotype on released mucin glycans compared to their glycoproteins (**Figure 1A-D**).

## Discussion

A systematic analysis of human gut bacteria that can grow on mucins or their components remains to be conducted. However, the data we present here underscore the idea that multiple sources of mucin (gastric vs. colonic) as well as forms (intact mucin domains or released, free *O*-glycans) should be considered to enhance knowledge of how bacteria access nutrition from complex mucin substrates. Previous reports have connected the activity of mucin-degrading bacteria to mucus defects *in vivo* and subsequent disease development^6–10^. Thus, it is imperative to understand the mechanisms by which different mucin-degraders catabolize these highly complex and sterically hindered mucins as well as the implications of how certain mucin-degraders cooperate to utilize (such as *R. torques* and *B. thetaiotaomicron*) or compete for (such as *R. torques* and *A. muciniphila*) this important substrate. Here, we advance the understanding of the mucin glycan-degrading mechanisms deployed by *R. torques*. Of note, *R. torques* utilizes both mucin glycoprotein domains and released oligosaccharide substrates from the stomach and the colon, degrades these substrates largely with secreted enzymes, and releases products from intact gastric mucins that become available to *B. thetaiotaomicron*. These findings establish *R. torques* as a potential keystone mucin-degrader which might be a possible target to block mucus barrier failure *in vivo,* the latter being an especially important point given this species’ emerging associations with IBD.

The mucin-degrading abilities of *R. torques* illuminate interesting differences between it and other known mucin-degrading species, which also fosters understanding of its role within the context of the gut community. First, its general lack of regulation of genes encoding putative secreted CAZymes in response to the presence of *O-*glycans differs from what has been observed in species like *B. thetaiotaomicron* and other *Bacteroides*^6,27,44^. While *R. torques* degradation of intact mucin glycoprotein domains was inhibited by prior growth on released free oligosaccharides (gMO), exposure to gMO strongly induces many loci encoding mucin-degrading enzymes in *B. thetaiotaomicron*^5,52,53^. The ability of *R. torques* to readily degrade intact mucin glycoproteins after growth on glucose, including the supernatant from these cultures, suggests that these enzymes are constitutively expressed during growth in non-mucin substrates, likely reflecting its adaptation to utilize mucins and mucin components as a nutrient source. Despite a lack of transcriptionally downregulated CAZymes during growth on gMOs, these enzymatic activities are repressed in the presence of gMO, which suggests a potential negative feedback loop in which released glycans from mucin glycoproteins suppress these systems in *R. torques* through post-transcriptional mechanisms. Such a system in *R. torques* may be fortuitously protective for the host by preventing excessive mucus erosion when free *O-* glycans are in excess. In cases where *R. torques* is associated with onset or exacerbation of disease, disruption of this observed repression could be a contributing factor.

Identification of putative mucin-degrading enzymes in *R. torques* supernatant revealed that many of these enzymes are multi-domain and large (17/18 of highly expressed CAZymes are >125 kDa), which poses a barrier to performing studies to recombinantly express and test activities of individual enzymes. However, investigations of the products generated by *R. torques* or structures recalcitrant to degradation, paired with cross-feeding experiments, nevertheless allowed us to gain mechanistic insight into the mucin-degradation strategy of *R. torques* and its implications for the gut community. The observation that *R. torques* supernatants contain sufficient enzyme activity to degrade mucin glycoproteins, suggests a predominantly extracellular degradation mechanism. This strategy stands in contrast to other known mucin-degraders. Namely, *A. muciniphila* is thought to import mucin glycoproteins or their fragments and degrade them intracellularly^41^.

Since *R. torques* likely uses extracellular degradation mechanisms, both released products and recalcitrant, incompletely degraded structures would become accessible to other species within the community. Our experiments suggest that *R. torques* releases monosaccharides and oligosaccharides from intact mucin glycoprotein but leaves the majority of sulfated glycans attached to the polypeptide backbone. By generating new products from these complex mucin glycoprotein substrates, *R. torques* generates a new pool of mucin-derived substrates available to other species, including those that are unable to access portions of intact mucin glycoprotein. Thus, these findings support the role of *R. torques* as a potential keystone mucin-degrader that likely impacts other species within the gut community.

The inner colon mucus layer largely acts as a filter and excludes particles the size of bacteria^16,54,55^. However, smaller particles are readily passed through this inner mucus layer, especially in the regions between the crypt openings^56^. While the molecular mechanisms of its extensive repertoire of mucin-degrading enzymes is still being elucidated, *B. thetaiotaomicron* is thought to predominantly employ outer membrane-bound or periplasmic enzymes for this degradation^31^. This suggests the use of a selfish mechanism, whereby *B. thetaiotaomicron* retains mucin byproducts for its own metabolism, thus not cross-feeding to other species and performing mucin glycan degradation outside of the inner protective mucus layer. In contrast, our data suggest that *R. torques* secretes many of its mucin glycan-degrading enzymes, and although they are larger than typical glycoside hydrolases, they may be able to diffuse into the inner mucus layer. Thus, *R. torques* may commence mucin degradation and lower the protective capacity of the inner mucus layer, which is one potential reason for its emerging association with IBD. These effects may be further exacerbated *in vivo* in the context of a more robust and diverse microbial community, which may provide additional critical enzymatic activities that *R. torques* is lacking, such as sulfatases, that may work cooperatively to enhance the degree of mucin degradation. Understanding the role that *R. torques* plays as a primary mucin-degrader may provide critical insight into the impairment of the colon mucus layer as a potential first step in the progression of spontaneous inflammation in the colon. Better understanding of the molecular mechanisms of bacterial mucin degradation will also open potential avenues for development of novel therapeutic targets to block IBD development.

## Materials and Methods

### Preparation of mucin substrates

Porcine gastric mucin (PGM) was purified as previously described^24^. Briefly, porcine gastric mucus (American Laboratories) was dissolved in 0.1 M NaCl and 0.02 M phosphate buffer at pH 7.8, stirred for an hour, then adjusted to pH 7.4 and stirred overnight. The PGM was centrifuged (10,000xg at 4°C for 30 minutes), the supernatant was collected and precipitated with cold ethanol (60% v/v). The precipitate was isolated by centrifugation (10,000xg at 4°C for 30 minutes). The ethanol precipitation was repeated once again, and the precipitate was dialyzed against MilliQ water and subsequently lyophilized.

Gastric mucin *O*-glycans (gMO) were generated from porcine gastric mucin (Type III, Sigma), adapted from a previously described method^57^. Briefly, PGM was autoclaved for 5 minutes in 100 mM Tris pH 7.4 and cooled to 65C. Proteinase K was added, and the solution was incubated overnight at 65°C. The solution was then centrifuged (8,000rpm for 30 mins). The supernatant was retained, and 0.4M NaOH and 1M NaBH_4_ were added. The solution was maintained at 65°C overnight and then neutralized to pH 7. The solution was again centrifuged and subsequently filtered through a 0.22 µm filter. To concentrate the solution, it was lyophilized, resuspended in ∼1/10 original volume in MilliQ water, dialyzed against MilliQ water, before being lyophilized again. The lyophilized powder was resuspended in 50 mM Tris pH 7.4 and passed over a DEAE column twice to remove contaminating glycosaminoglycans (flow through from anion exchange column was retained as the neutral gMO fraction).

Colonic mucin glycoprotein (cMUC2) was purified from porcine colons (anus through ∼2 feet into the colon) obtained from the Michigan State University Meat Laboratory. cMUC2 was purified using previously reported procedures^31^. Briefly, the mucosa was scraped from the colons and stored at −80°C until purified. Mucosal scrapings were homogenized into extraction buffer (6 M GuHCl, 5 mM EDTA, 0.01 M NaH2PO4; pH 6.5). The mucosal scrapings stirred in extraction overnight at 4°C, then were pelleted by centrifugation (18,000rpm for 30 mins at 4°C). This extraction process was repeated six times. After the sixth extraction, the pelleted mucin was resuspended in reduction buffer (6 M GuHCl, 0.1 M Tris, 5 mM EDTA; pH 8.0) and 25 mM dithiothreitol was added before shaking the solution for 5 hours at 37°C. Then iodoacetamide was added in a ratio of 2.5 times the molar amount of added dithiothreitol, and the solution was stirred overnight at room temperature. The solution was then centrifuged (10,000rpm for 30 mins at 4°C) and the soluble mucin was collected, dialyzed against MilliQ water, and lyophilized. The mucin was then resuspended in trypsin digestion buffer (0.1 M Tris pH 8.0). Trypsin was added to a concentration of 1 mg/mL and the solution was incubated overnight at 37°C. An additional 0.5 mg/mL of trypsin was added the next day, the solution was incubated for 5 hours at 37°C, and was boiled and cooled before dialyzing against MilliQ water and lyophilizing. Colonic mucin oligosaccharides (cMO) were purified from cMUC2 using the same method as used to generate gMO, except they were not run over the DEAE column due to the high abundance of sulfated glycans in colonic mucus.

### Bacterial strains and mucin growth curves

All bacterial cultures were grown in an anaerobic chamber (Coy Labs; 85% N_2_, 10% H_2_, 5% CO_2_) at 37°C. *R. torques* VIII-239, a previously described human isolate, was grown overnight in a previously described modified version of the common YCFA medium^37,58^. *B. thetaiotaomicron tdk*^-/-^ strain (a thymidine kinase-deficient mutant in *B. thetaiotaomicron* VPI-5482 used to generate allelic exchange mutants^50^) was used throughout this study and was grown overnight in TYG medium^59^ and *A. muciniphila* DMS 22959 was grown overnight in a custom chopped meat medium^49^. To evaluate growth on mucin derivatives, cells from rich medium culture were pelleted, washed in respective carbohydrate free media (*Rt:* YCFA no glucose, *Am:* chopped meat no sugars; *Bt: Bacteroides* minimal media no glucose^27^ and diluted (1:40) before 100 µL of washed cells were added to 100 µL of growth substrates in 96 well plates (Costar). Growth substrates were tested at the following concentrations: glucose/N-acetylglucosamine (5 mg/mL), mucin *O*-glycans (gMO, 10 mg/mL), porcine gastric mucin (PGM, 10 mg/mL), colonic *O*-glycans (cMO, 5 mg/mL), colonic mucin glycoprotein (cMUC2, 2.5 mg/mL). Growth was measured by optical density at 600nm (OD_600_) at ∼10-minute intervals using a microplate stacker (BioTek BioStack) and microplate reader (BioTek PowerWave). Growth curve OD_600_ values were normalized by subtracting average baseline readings (from the first three OD readings) from all readings to correct for the background OD_600_ of growth substrates.

### Gel electrophoresis and Periodic acid Schiff staining

*R. torques* cultures were grown overnight in YCFA with glucose (5 mg/mL) or gMO (10 mg/mL). Culture samples were taken and added to reaction mixtures or supernatants were collected by centrifugation for 2 minutes at 8,000rpm prior to addition to reaction mixtures. Culture or supernatant samples were added 1:1 to cMUC2 (2.5 mg/mL) or PGM (10 mg/mL) and were incubated anaerobically for 2 days. Samples were collected, diluted 1:4 in Laemmli sample buffer (Bio-Rad) and heated for 10 minutes at 98°C. Samples were loaded onto 4-12% Bis-Tris gradient gels (Thermo Fisher) (with the exception of the filtered vs. spun down supernatant PAS gel, which was run on a 4-12% Tris-Glycine gel (Thermo Fisher) and run for 60 mins at 180V. Gels were stained as previously described^60^. Briefly, gels were incubated with 50% methanol for 30 minutes and washed with 5% acetic acid 3 times for 10 minutes each. Gels were incubated in 0.7% periodic acid in 5% acetic acid for 3 hours, followed by 3 washes in 5% acetic acid for 5 minutes each. Lastly, gels were incubated overnight in Schiff’s reagent (Fisher) and imaged using Bio-Rad GelDoc Imaging System. Intensity of mucin bands were quantified by densitometry with ImageJ, and intensity from a blank lane in each gel was subtracted from all other lanes in the respective gel to control for background staining across gels. The percent degradation in various conditions was calculated relative to controls lacking *R. torques* culture or supernatant.

### RNA-sequencing of R. torques on glucose vs. gMO

*R. torques* overnight cultures in YCFA with glucose were pelleted, washed in YCFA with no carbon source, and resuspended. *R. torques* was then diluted 1:40, and 100 µL of washed culture was added to 100 µL of substrate (5 mg/mL glucose or 10 mg/mL gMO) in a 96 well plate (Costar). Washed culture from each overnight replicate was added to the plate in 12 replicate wells to increase biomass for RNA extraction. Growth was monitored by OD_600_ using a microplate stacker and plate reader. Cultures were collected during mid-log phase (OD_600_ ∼0.7-0.9) and the 12 replicate wells were pooled to form a single sample. These were harvested, treated with 1 mL of RNAprotect Bacteria Reagent (Qiagen), and pellets were stored at −80°C until RNA isolation. Samples were resuspended in 500 µL Buffer A (200 mM NaCl, 200 mM Tris, 20 mM EDTA, pH 8), 210 µL 20% SDS, and 500 µL phenol:chloroform:isoamyl alcohol (pH 4.5) and bead beat on high for 5 minutes (Minibeadbeater, Biospec Products). Samples were centrifuged at 18,000xg for 3 minutes at 4°C, and the aqueous phase was moved to a new tube. 500 µL phenol:chloroform:isoamyl alcohol was added and mixed, and samples were centrifuged again at 18,000xg for 3 minutes. The aqueous phase was moved to a new tube, and 50 µL of 3M sodium acetate and 500 µL of cold isopropanol was added. Samples were stored at −80°C for at least 20 minutes and centrifuged for 20 minutes at 18,000xg and 4°C. The pellet was washed with 500 µL of cold 70% isopropanol and centrifuged again at 18,000xg for 5 minutes. The pellet was allowed to dry and was resuspended in 50 µL of water. The samples were then purified using the RNeasy Mini Kit (Qiagen) using Protocol 1: Enzymatic Lysis of Bacteria, then Protocol 7: Purification of Total RNA from Bacterial Lysate Using the RNeasy Mini Kit. Eluted RNA was treated with DNase (NEB) in the following reaction mixture: 4 µL DNase, 10 µL 10X DNase buffer, 50 µL RNA, 36 µL water. Reactions were incubated for 30 minutes at 37°C, then inactivated with 4 µL of 0.5M EDTA. Samples were again purified using the RNeasy Mini Kit. rRNA was depleted from the samples using the MICROBExpress Bacterial mRNA Enrichment Kit (ThermoFisher). RNA was submitted to the University of Michigan Advanced Genomics Core for library preparation and RNA-sequencing (Illumina NovaSeq platform, 150 bp read length, paired end).

Reads were analyzed using previously reported methodology^48^. Briefly, Trimmomatic v0.39 was used to filter reads for quality. Reads were aligned to the *R. torques* genome using BowTie2 v2.3.5.1. Reads mapping to gene features were counted using htseq-count (release_0.11.1). Differential expression analysis was performed with the edgeR v3.34.0 package in R v.4.0.2 (with the aid of Rstudio v1.3.1093). Library normalization was performed using the trimmed mean of M-values (TMM) method. Integrated Genome Viewer (IGV) was used to visualize coverage data.

### Proteomics

*R. torques* overnight cultures in YCFA with glucose were grown and centrifuged at 7,830rpm for 2 mins. Samples were filtered where noted using a 0.22 μM PVDF syringe filter. Proteins were precipitated using a pyrogallol red methanol molybdate (PRMM) solution. PRMM was made with 20% methanol, 0.05 mM pyrogallol red, 0.16 mM sodium molybdate, 1 mM sodium oxalate, 50 mM succinic acid and pH adjusted to pH 2. PRMM was added 1:1 with isolated supernatants, and pH was adjusted to pH of 3. Samples were incubated for 2 hours at room temperature and then overnight at 4°C. Samples were then centrifuged for 60 mins at 10,000xg at 4°C and pellets were washed with cold acetone. Centrifugation and washing were repeated once, and pellets were dried and resuspended in 8 M urea with 50 mM HEPES at pH 8.

Samples were submitted to the Proteomics Resource Facility (PRF) and were analyzed using high-resolution LC-MS/MS analysis per the protocol optimized by the PRF and previously described^48^. The following modifications were made from the previous protocol: cysteines were alkylated with 65 mM 2-Chloroacetamide, trypsin digestion was performed after diluting urea to <1.2 M and using 0.5 µg trypsin (Promega), and processed peptides were dissolved in 20 µl of buffer A. Peptides were resolved over a period of 90 min (2-25% buffer B in 45 min, 25-40% in 5 min, 40-90% in 5 min, then held at 90% buffer B for 5 min and equilibration with buffer A for 30 min). Proteins were identified by searching the generated MS/MS data against a database of all protein sequences present in *R. torques* VIII-239. Parameters used were the same as previous, except for the use of a 0.1 kDa fragment tolerance.

### R. torques supernatant enzyme activity assays on model glycans

*R. torques* was grown in large batch cultures (40 mL) overnight in YCFA with glucose. Cultures were then spun at 7,200xg for 10 mins, which was repeated as necessary until cells were pelleted. Supernatants were decanted and 60 mL of 4 M ammonium sulfate was added. Samples were incubated for 30 mins at room temperature and then centrifuged at 18,000xg for 30 mins at 4°C. The supernatant was decanted, and the precipitate was resuspended in 1 mL of PBS. Samples were dialyzed overnight into PBS in 3.5 kDa cutoff dialysis tubing at 4°C. Reaction mixtures of the following final composition were made: 2-5 mM substrate, 50 mM phosphate buffer pH 7, and 10 µL resuspended ammonium sulfate precipitated proteins. Final substrate concentrations were 2 mM (Lewis a,b,x,y; blood groups A and B) or 5 mM (LacNAc and LNB). Lewis antigens and sulfated sugars were sourced from Dextra, lacto-*N*-biose and *N-* acetyllactosamine were sourced from CarboSynth/Biosynth, and blood group A and B were sourced from Elicityl. Reactions testing for activity on sulfated sugars were made of the same composition, except for the use of 10 mM MES buffer with 5 mM CaCl_2_ pH 6.5. *B. thetaiotaomicron* was grown overnight in Bacteroides minimal medium with 5 mg/ml final concentration of glucose, cells were collected, washed, and resuspended in Bacteroides minimal medium with 5 mg/ml keratan sulfate and incubated for one hour prior to sonication (30% amplitude, 1s on 9s off, 30 seconds total). A 10 ul sample of this preparation was added to the same reaction composition as a positive control for sulfatase activity. Reactions were incubated overnight at 37°C and stored at −20°C until analysis.

Samples were diluted 1:10 in water and were analyzed by High Performance Anion Exchange Chromatography with Pulsed Amperometric Detection (HPAEC-PAD, Dionex ICS-6000 HPIC System, Thermo Scientific) using a CarboPac PA100 column (Thermo Scientific). The following buffers were used: Buffer A (100 mM NaOH), Buffer B (100 mM NaOH, 500 mM sodium acetate), Buffer C (water). The following method sequence was utilized for all samples with a constant flow rate of 1 mL/min: 15 minute pre-equilibration with 90% Buffer C, 10% Buffer A; 20 min 90% Buffer C, 10% Buffer A; 20 min gradient from 90% Buffer C, 10% Buffer A to 100% Buffer A; 20 min gradient from 100% Buffer A to 100% Buffer B; 10 min 100% Buffer B; 10 min 90% Buffer C, 10% Buffer A). Chromatograms were analyzed and peaks were quantified in Chromeleon Chromatography Data System Software (ThermoFisher) using the Basic Quantification method. Standard curves of fucose, galactose, and *N*-acetylglucosamine were created to calculate concentrations of respective sugars from peak areas.

Reactions to assess activities on sulfated glycans were prepared in the same manner as for the blood group and Lewis antigens. Reactions were analyzed using thin layer chromatography (TLC). Each reaction was spotted (3 µL) onto a silica plate and was placed in 100 mL of running buffer of butanol:water:acetic acid (2:1:1). Once the solvent front reached the end of the plate, the plate was dried and submerged in developer (2 g diphenylamine, 100 mL ethyl acetate, 2 mL aniline, 10 mL 85% H3PO3, 1 mL 37.5% HCl) for ∼30 seconds, and were dried again. Sugars were visualized by heating the plate over a flame.

### Analysis of released monosaccharides from mucin glycoprotein substrates

*R. torques* cultures were grown overnight in YCFA with glucose, and aliquots were removed and centrifuged at 8,000rpm for 2 mins. Supernatants were isolated and incubated with mucin substrates PGM (10 mg/mL final concentration) or cMUC2 (2.5 mg/mL final concentration) 1:1 by volume. Samples were incubated anaerobically at 37°C and were stored immediately (0h control) or incubated for 24h. Samples were analyzed using the L-Arabinose/D-Galactose Assay Kit (Megazyme) and the L-Fucose Assay Kit (Megazyme) per kit protocols for the microplate assay format. Baseline absorbance for each sample and the absorbance from the corresponding negative control were subtracted from sample absorbance readings and concentrations were calculated from a standard curve for each corresponding monosaccharide.

### Bioinformatic analysis of putative mucin-degrading enzymes

The presence of potential signal sequences to indicate secreted proteins in *R. torques* for transcriptomic/proteomic analyses were identified using SignalP 5.0 (DTU Health Tech). Proteins identified as top hits from transcriptomic or proteomic datasets lacking a functional annotation (*i.e.*, annotated as hypothetical proteins) were analyzed for putative functions using NCBI Protein BLAST (BlastP). Domain analysis of putative mucus-degrading enzymes was performed using InterPro scan by sequence, using default search settings. Domains predicted as F5/8 type C domains are annotated as putative CBM domains. *R. torques* VIII-239 amino acid sequences were analyzed using SulfAtlas HMM using default settings and cutoffs (high prediction: score>300; low prediction: 200 < score < 300 and coverage > 80%; probable fragment: 100 < score < 300 and coverage < 80%)^61^.

### LC-MS/MS analysis of mucin glycans before and after R. torques growth

*R. torques* was grown on cMUC2 and PGM using the same growth methods described above, and samples were collected immediately after inoculating cultures (pre-growth) and after cultures reached stationary phase (post-growth). Glycans from the samples were isolated by dot blot using a PVDF membrane and were analyzed using carbon-column liquid chromatograph-electrospray ionization tandem mass spectrometry (LC-ESI/MS), as previously described^62,63^.

### Growth of bacterial strains on R. torques supernatant digests of PGM

*R. torques* was grown overnight in YCFA with glucose, and then was washed in YCFA with no additional carbon source and diluted 1:80 in YCFA with PGM (10 mg/mL) as the main carbon source. One sample was collected immediately to serve as a control (0h) and the other sample was incubated anaerobically at 37°C for 24h. Upon collection, samples were boiled for 10 minutes and then centrifuged at 7,197xg for 10 minutes. The supernatant was collected and dialyzed (1KDa cutoff) against 4 L water, changing the water three times. Samples were then lyophilized, resuspended at 20 mg/mL in PBS, and autoclaved with a 5-minute sterilization cycle time. Growth experiments were performed as previously described, except strains were grown in partially defined medium (DM)^48,49^.

### Co-culture of R. torques and B. thetaiotaomicron on PGM

*R. torques* was grown overnight in YCFA with glucose and *B. thetaiotaomicron* was grown overnight in TYG. A 1 mL sample was taken from each culture, centrifuged for 2 mins at 8,000 rpm and was washed in 2X DM. Centrifugation was repeated and cultures were resuspended in 2X DM. Each culture was added in a 1:1 volume ratio (10 µL each species), except in one experiment, where they were added in a 3:1 ratio (15 µL *R. torques* and 5 µL *B. thetaiotaomicron)* to a 96 well plate pre-loaded with 80 µL of 2X DM and 100 µL of substrate in a 96 well plate (5 mg/mL final concentration of glucose or 10 mg/mL final concentration of PGM). Co-cultures were grown anaerobically at 37°C and were passaged every 24 hours. Samples were collected and frozen at −20°C until DNA was extracted using the DNeasy Blood and Tissue Kit (Qiagen) according to the Protocol: Pretreatment for Gram-Positive Bacteria, followed by the Protocol: Purification of Total DNA from Animal Tissues Spin-Column as described. DNA was quantified using a NanoDrop spectrophotometer (Thermo Scientific) and diluted to 5 ng/µL. qPCR reactions were performed as previously described^48^, using *B. thetaiotaomicron* (BT1927F: 5-TCGCCTTTTTCAGATCAGTAGTTGG-3; BT1927R: 5-ACGAAAATGGAGTTGAATGGAATAAGTT-3) or *R. torques* (RT1159F: 5-CCGTTCCTGACAACATTAG-3; RT1159R: 5-GCC GAT TCT CTT TCC TAT ATC-3) specific primers with corresponding standard curves generated from purified genomic DNA from each respective species.

### Data Availability

RNA-sequencing data is deposited to the NIH Sequence Read Archive under accession number (pending). LC-MS/MS data from the *R. torques* proteomics experiment is deposited in the Proteomics Identification Database (PRIDE) under accession number (pending). Raw LC-MS/MS data from mucin glycans before and after growth of *R. torques* is deposited in GlycoPOST under accession number (pending).

## Acknowledgements

We would like to acknowledge support from the University of Michigan Biomedical Research Core Facilities (Proteomics Resource Facility), which is supported by an NIH Center Grant (University of Michigan Center for Gastrointestinal Research, P30 DK034933). SRS was supported by funds from the NIH T32 Cellular Biotechnology Training Program (T32 GM145304). We are extremely grateful for support from the Kenneth Rainin Foundation (Innovator Award to ECM) and the US National Institutes of Health (R01s DK118024, DK125445 to ECM). MPO was supported by funds from UL1TR002240. GCH was supported by funds from Knut and Alice Wallenberg Foundation (2017.0028), Swedish Research Council (2017-00958), the European Research Council ERC (101100663). We acknowledge the Proteomics Core Facility at the University of Gothenburg. We thank SciLifeLab and BioMS funded by the Swedish research council for providing financial support to the Proteomics Core Facility (CJ), Sahlgrenska Academy.

---

**Supplemental Figure S1.**
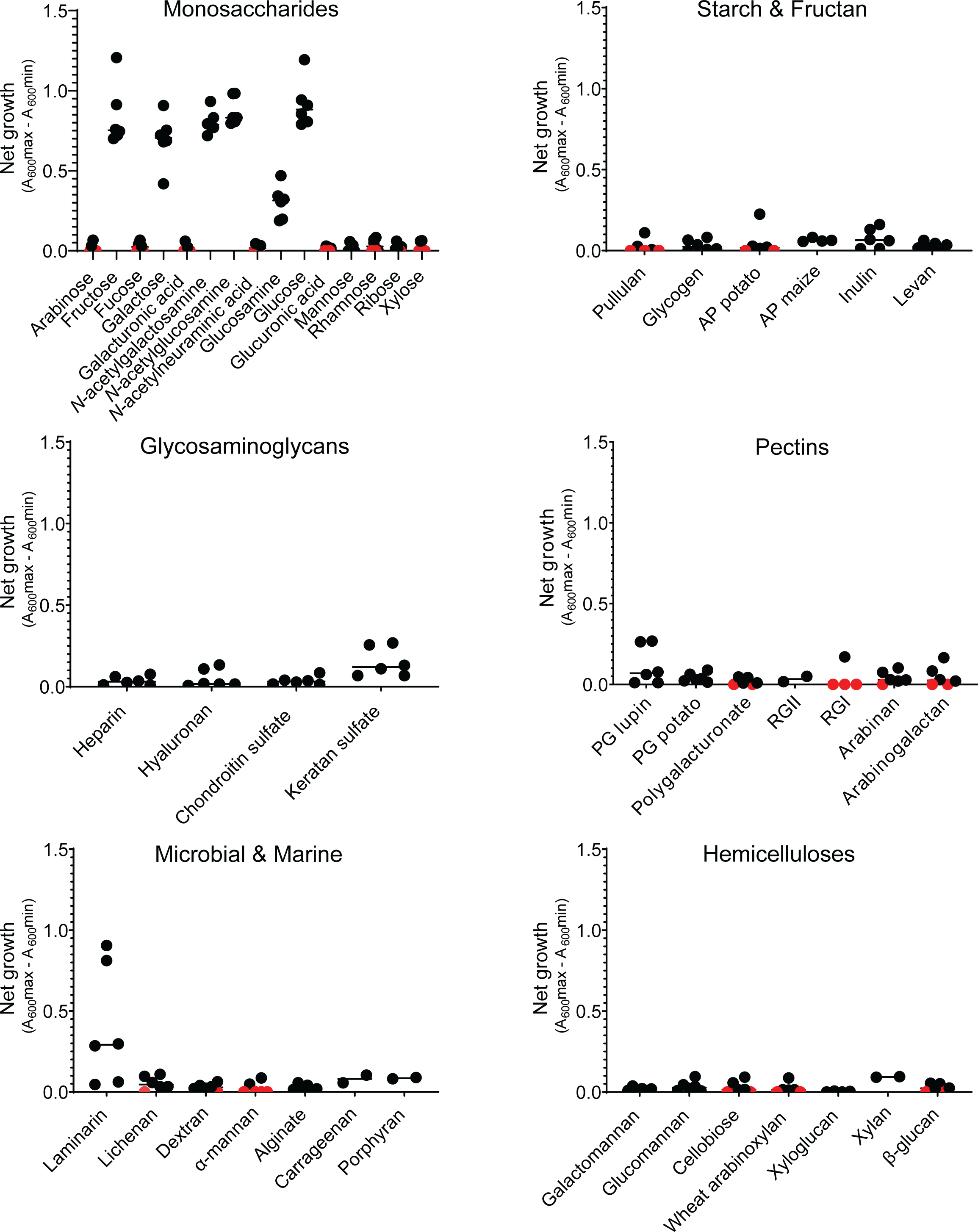
*R. torques* growth on various carbohydrates. *R. torques* was grown on 46substrates, each as the main carbohydrate source in YCFA medium. Growth was measured by absorbance at 600 nm for ∼92 hours. Growth curves were analyzed to identify minimum and maximum absorbance values for each curve (excluding values representing artifacts due to brief disturbances of the plates or baseline noise), and the net growth (max absorbance – min absorbance) was calculated. For growths in which the absorbance consistently decreased throughout the growth curve with no apparent increase in absorbance, net growth was reported as zero, and these points are displayed in red. AP = amylopectin, PG = pectic galactan, RG = rhamnogalacturonan (type I or type II specified).

**Supplemental Figure S2.**
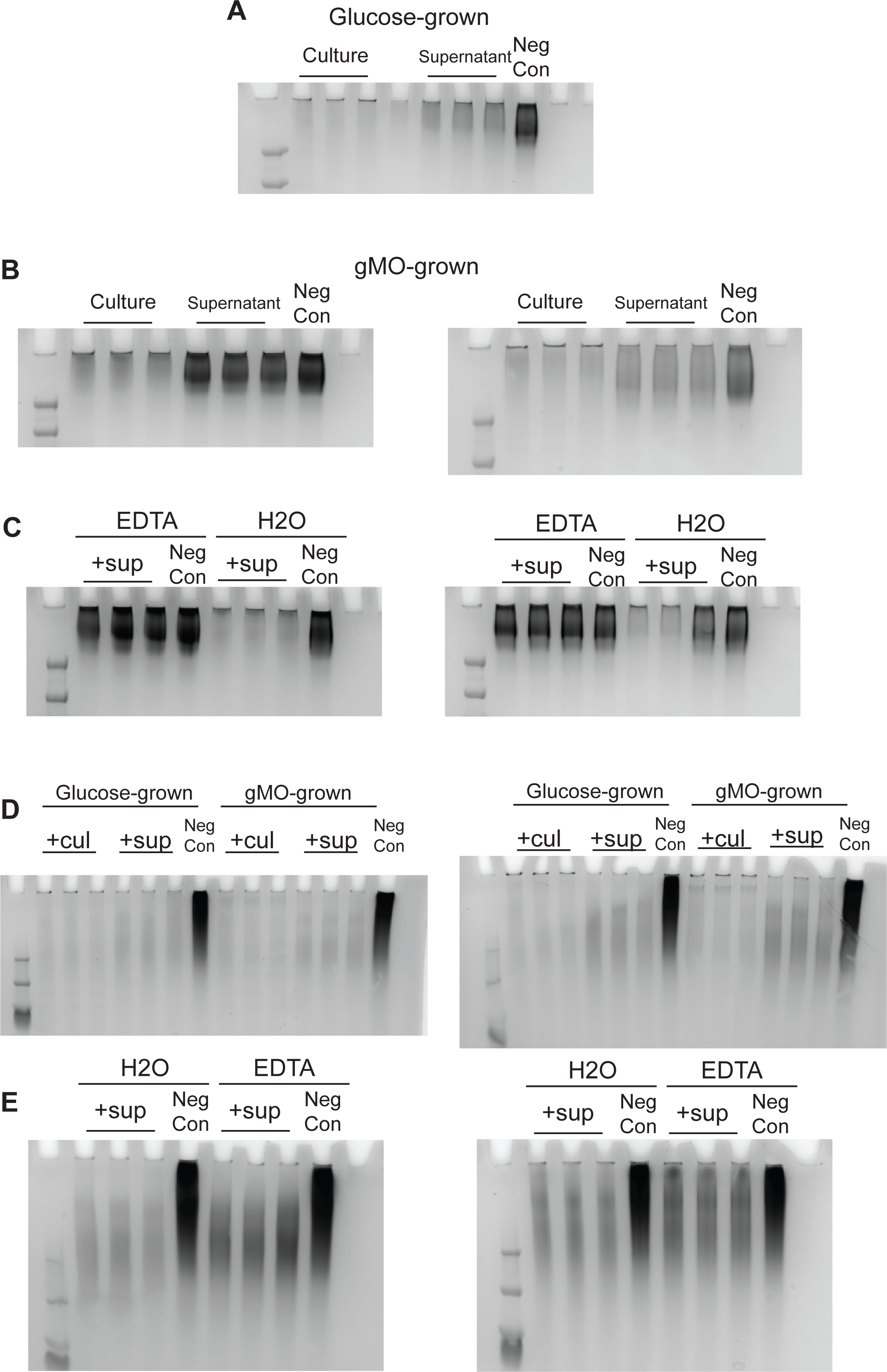
Periodic acid-Schiff-stained gels to visualize degradation of mucin glycoproteins by *R. torques.* Culture (*i.e*., cells and supernatants) or cell-free supernatant samples were collected from *R. torques* after overnight growth on glucose or gMO. Samples were incubated with cMUC2 or PGM anaerobically for 2 days and electrophoresed through 4-12% Bis-tris gels and stained with periodic acid-Schiff stain to visualize mucins. **A-C**, Degradation of cMUC2 (2.5 mg/ml final concentration) by glucose-grown samples (**A**), gMO-grown samples (**B**), or 10mM EDTA treated samples (**C**). **D-E**, degradation of PGM (10 mg/ml final concentration) by glucose or gMO-grown samples (**D**) or 10 mM EDTA treated samples (**E**).

**Supplemental Figure S3.**
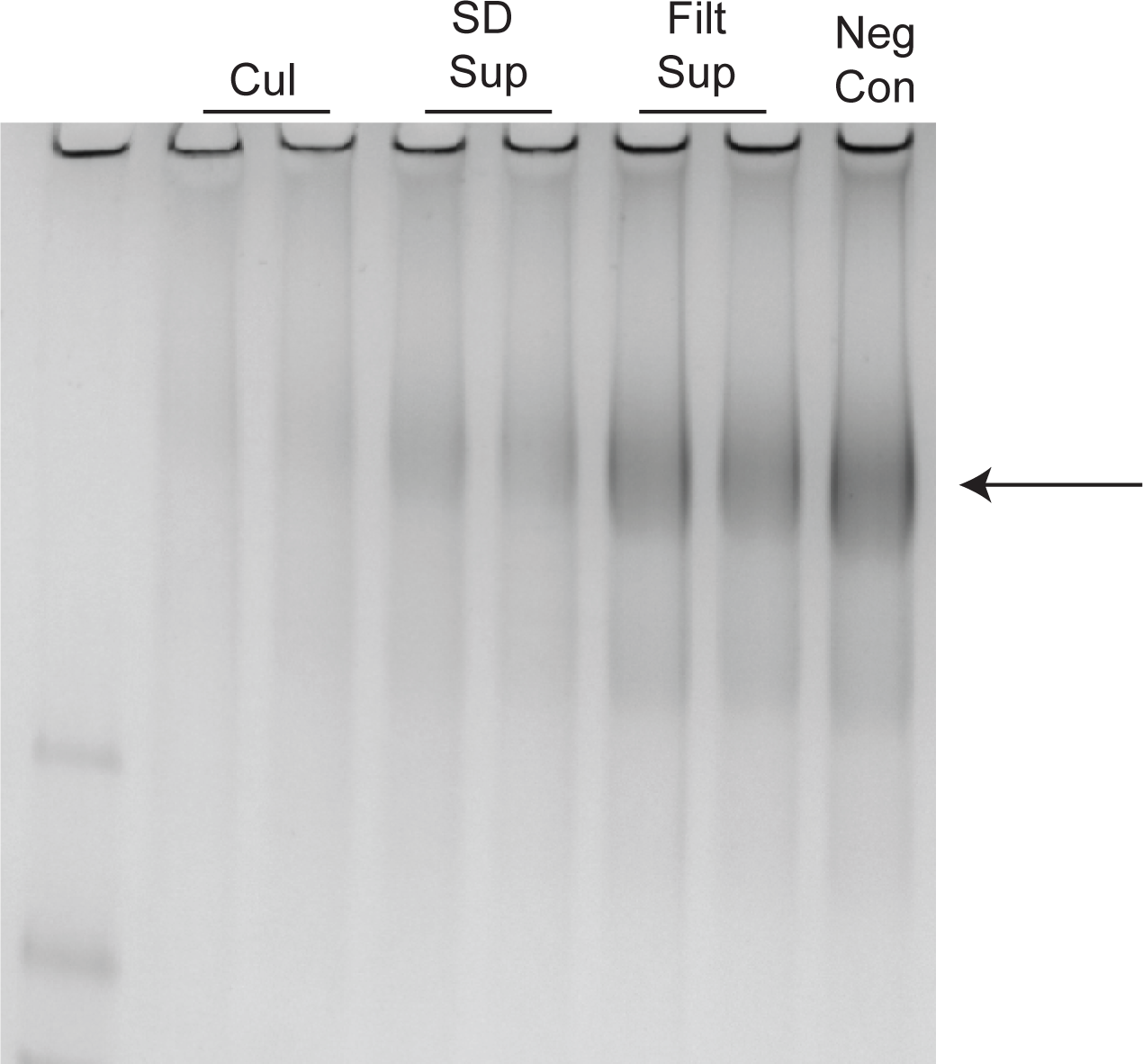
Degradation of rGP by *R. torques* supernatants after centrifugation or filtering. *R. torques* was grown overnight in YCFA with glucose and samples were either centrifuged or filtered (0.22 um PVDF filters) to collect supernatants. Supernatants were then incubated with rGP anaerobically for 2 days. Samples were analyzed by electrophoresis through a 4-12% Tris-Glycine gel and stained with periodic acid-Schiff stain to visualize mucin (major band indicated by arrow). Cul = whole culture (cells + supernatant), SD sup = spun-down (centrifuged) supernatant, Filt sup = filtered supernatant, Neg con = negative control (YCFA with glucose added in lieu of *R. torques* cultures or supernatant).

**Supplemental Figure S4.**
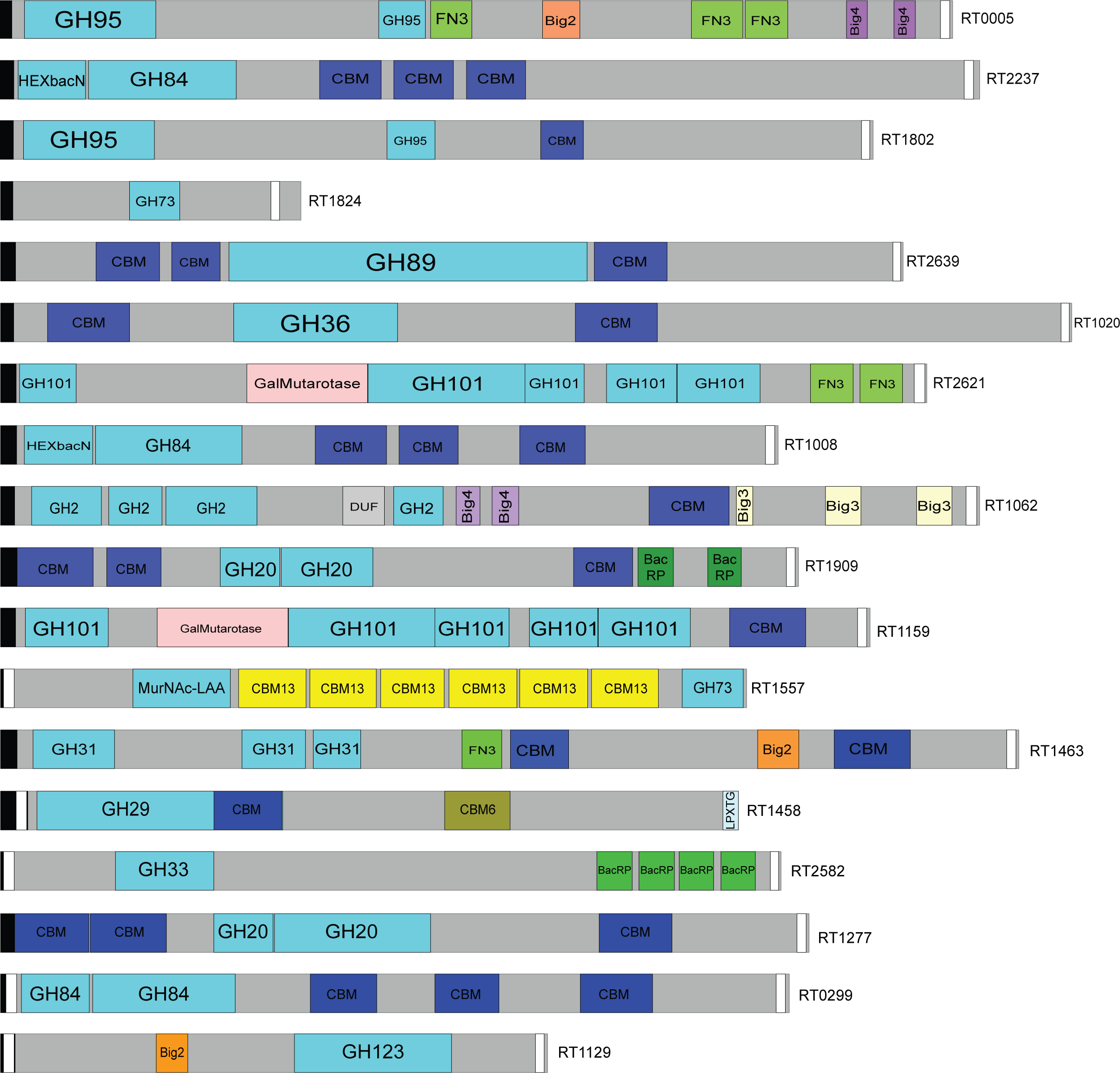
Predicted domain architecture of highly expressed and transcribed CAZymes. Domain architecture predictions were generated from Interpro 96.0 sequence searches. Predicted F5/8 type C domains are shown as putative carbohydrate binding module (CBM) domains. Figure made with Illustrator for Biological Sequences^64^. Black = signal peptide; white = transmembrane domain; teal = CAZyme domains; blue = CBM domains.

**Supplemental Figure S5.**
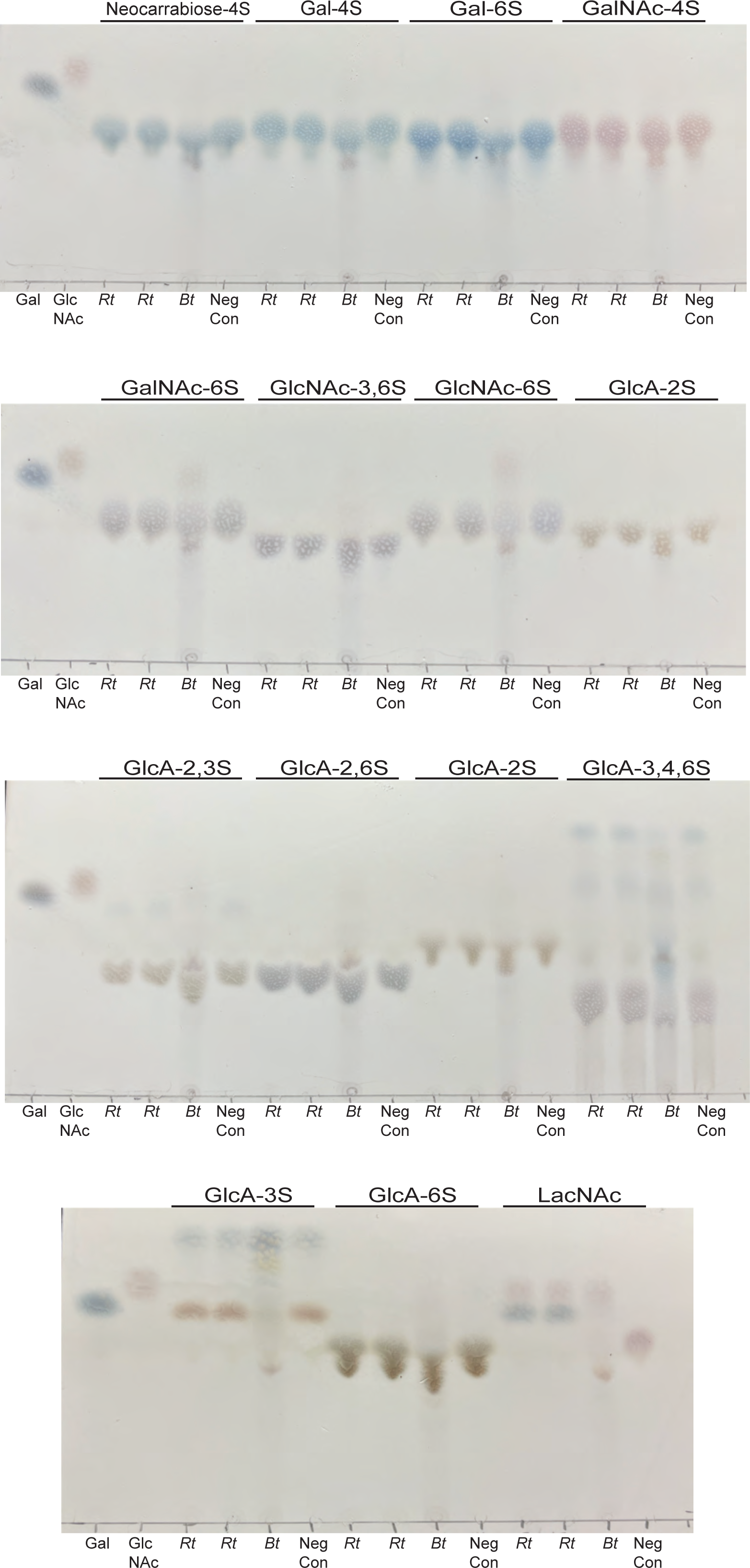
*R. torques* fails to degrade sulfated sugars. Proteins were ammonium sulfate precipitated from the supernatant of *R. torques* overnight cultures after growth in YCFA with glucose and incubated with respective sugars (5mM final concentration, except for GlcA-6-O-sulfate which was 20 mM final concentration). Reactions were analyzed by thin layer chromatography using diphenylamine-aniline-phosphoric acid developer. *Rt* indicates samples where *R. torques* precipitated proteins were added, *Bt* indicates samples where sonicated *B. thetaiotaomicron* cultures exposed to keratan sulfate was added, and Neg Con indicates negative control samples that contained buffer (10 mM MES and 5 mM CaCl_2_). Gal = galactose; GalNAc = *N*-acetylgalactosamine; GlcNAc = *N*-acetylglucosamine; GalA = galactosamine; GlcA = glucosamine.

**Supplemental Figure S6.**
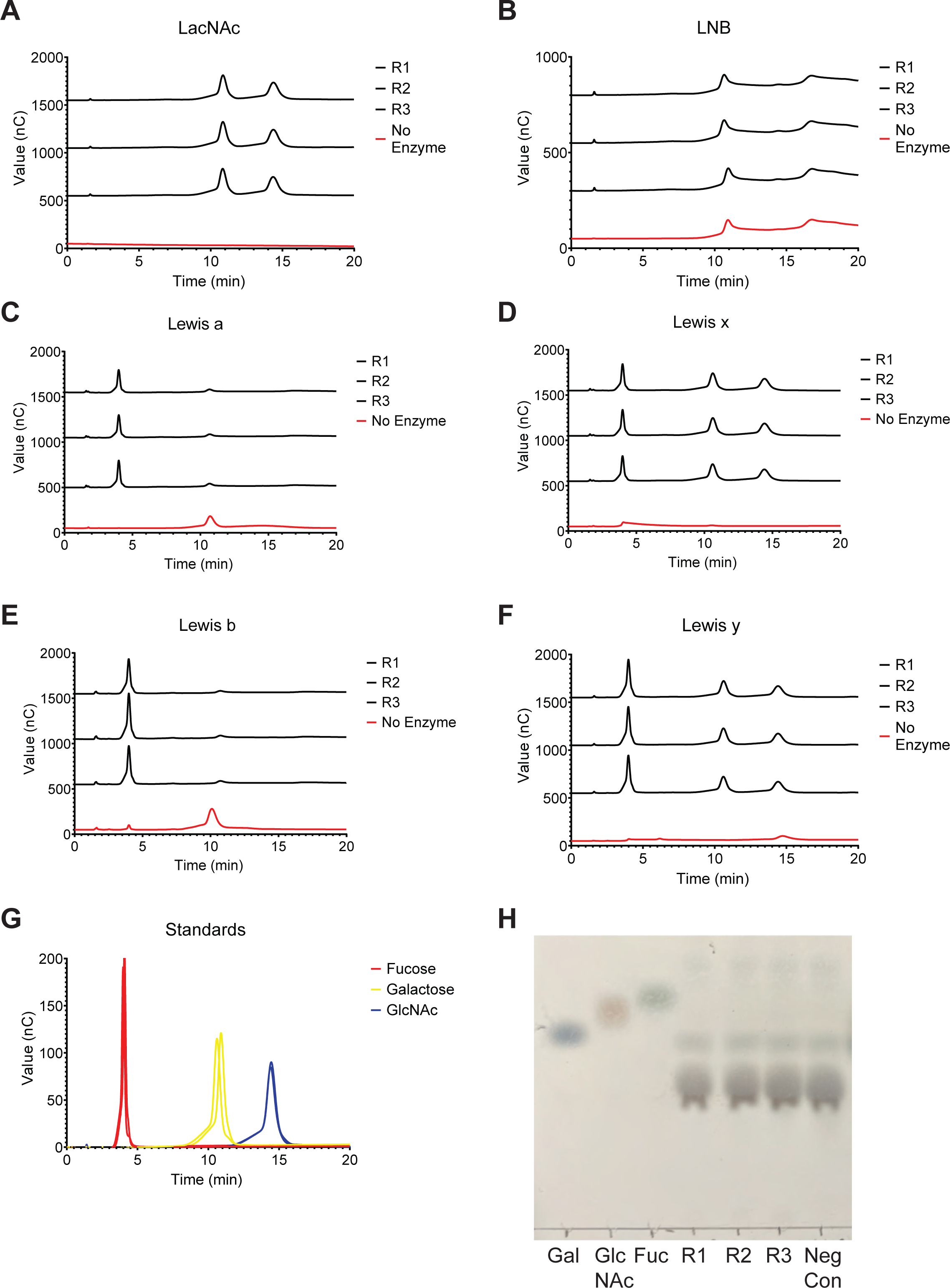
HPAEC-PAD analysis of *R. torques* ammonium sulfate precipitated protein activity on model mucin sugars. **A-G**, Chromatograms collected by HPAEC-PAD analysis of *R. torques* supernatant precipitated protein reactions with model glycans *N*-acetyllactosamine (LacNAc, **A**), and lacto-*N*-biose (LNB, **B**), Lewis a (**C**), x (**D**), b (**E**), y (**F**) compared to single monosaccharide standards (**G**). R1-R3 = replicates reactions with supernatant precipitates from 3 *R. torques* cultures. No enzyme control is shown in red, where PBS was added instead of *R. torques* precipitates. **H**, Thin layer chromatogram of reaction products of activity assay on lacto-*N*-biose (LNB) from A. R1-R3=replicate reactions with *R. torques* supernatant precipitates; Neg Con=no enzyme control reaction with PBS.

**Supplemental Figure S7.**
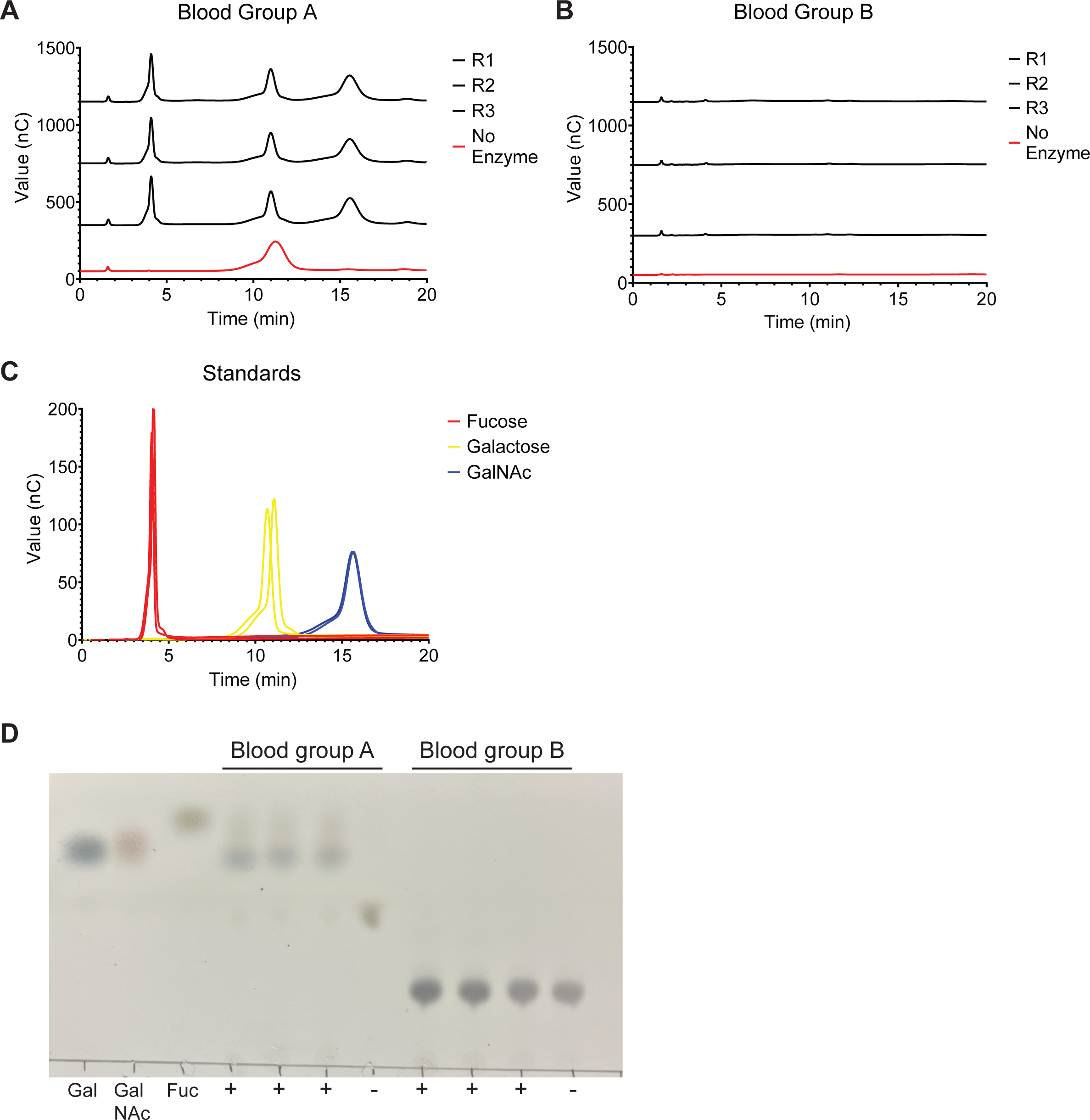
HPAEC-PAD analysis of *R. torques* ammonium sulfate precipitated protein activity on blood groups A and B. **A-C**, Chromatograms collected by HPAEC-PAD analysis of *R. torques* supernatant precipitated protein reactions with model glycans blood groups A (**A**) and B (**B**) compared to single monosaccharide standards (**C**). R1-R3 = replicates reactions with supernatant precipitates from 3 *R. torques* cultures. No enzyme control is shown in red, where PBS was added instead of *R. torques* precipitates. **D**, Thin layer chromatogram of blood group A and blood group B reactions from A. Gal=galactose, GlcNAc=*N*-acetylglucosamine, Fuc=fucose, R1-R3=replicate reactions with *R. torques* supernatant precipitates, Neg Con=no enzyme control reaction with PBS.

**Supplemental Figure S8.**
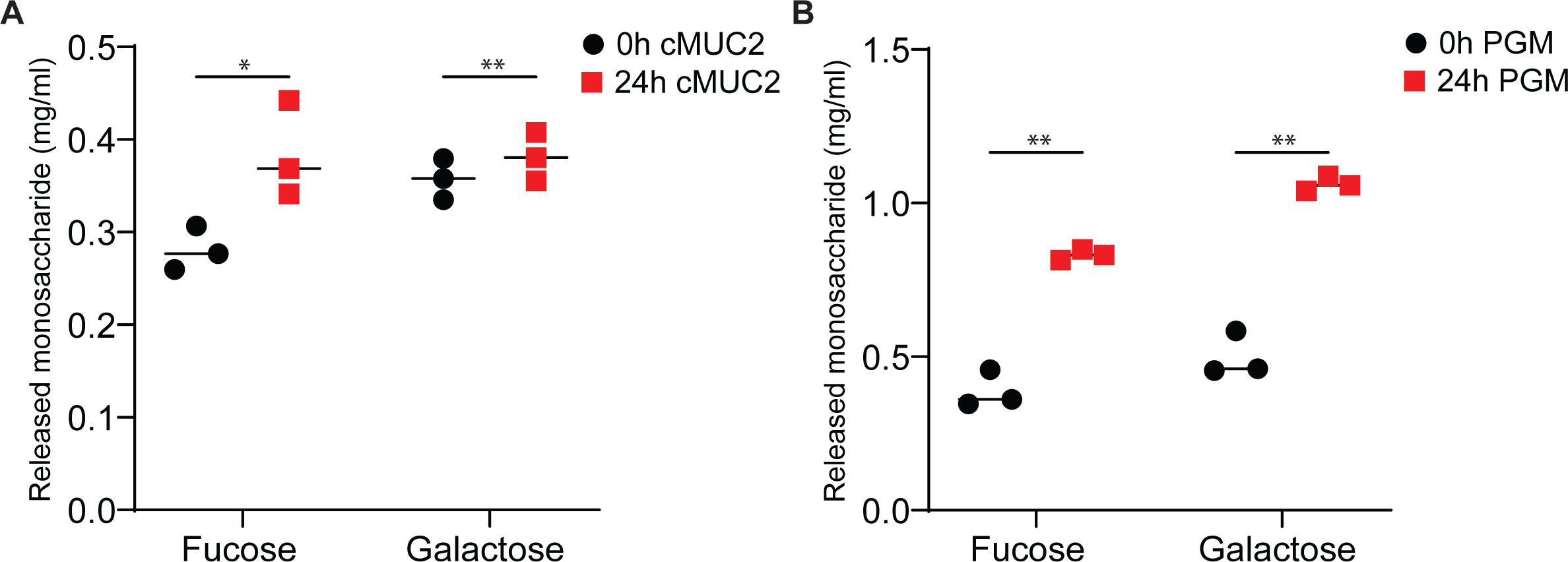
Fucose and galactose released from cMUC2 and PGM after digestion by supernatants from *R. torques* grown on gMO. **A-B**, Concentration of fucose or galactose released after 24-hour incubation of *R. torques* supernatants from cultures grown on gMO with cMUC2 (2.5 mg/ml final; **A**) or PGM (10 mg/ml final; **B**). Statistics were analyzed with paired, two-tailed t tests comparing release of each monosaccharide between substrates in each panel. *p<0.05, **p<0.01.

**Supplemental Figure S9.**
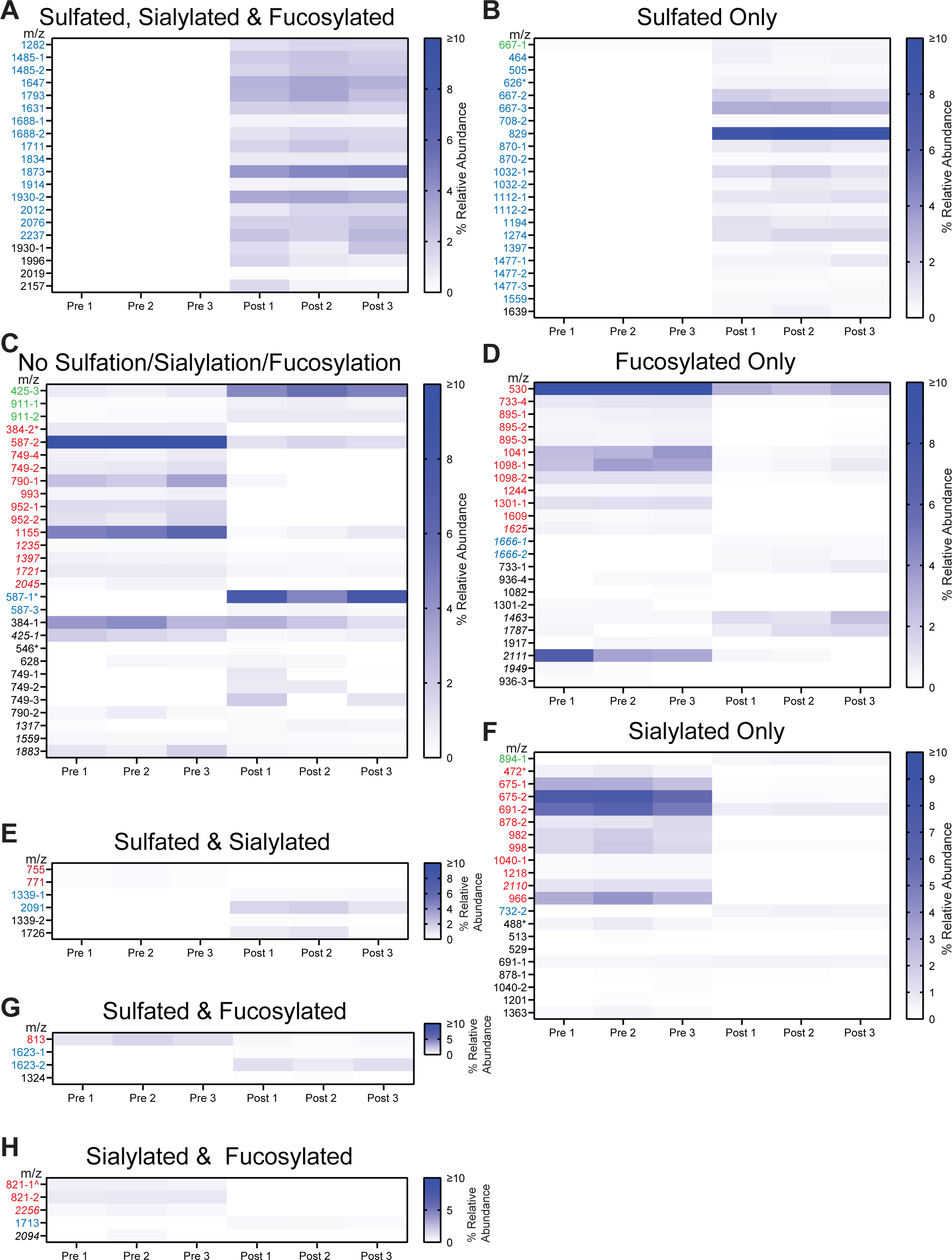
LC-MS/MS glycans identified before and after growth of *R. torques* on cMUC2. **A-H**, Glycans detected were grouped based on structural features: sulfated, sialylated and fucosylated (**A**), sulfated only (**B**), no sulfation, sialylation, fucosylation (**C**), fucosylated only (**D**), sulfated and sialylated (**E**), sialylated only (**F**), sulfated and fucosylated (**G**), and sialylated and fucosylated (**H**). Heat maps indicate percent relative abundance of each glycan pre- or post-growth of *R. torques.* M/z values in green represent glycans detected in at least one pre-growth sample that significantly increase in abundance post-growth; glycans in red decreased in abundance post-growth compared to pre-growth, glycans in blue were undetected pre-growth but were detected post-growth; and glycans in black were not significantly different in abundance pre- or post-growth. Statistics were analyzed using paired, two-tailed t-tests between the three pre- and three post-growth samples for each glycan. * = peeling reaction product, ^ = fuc-ol terminating glycan, italicized m/z values = *N*-glycan.

**Supplemental Figure S10.**
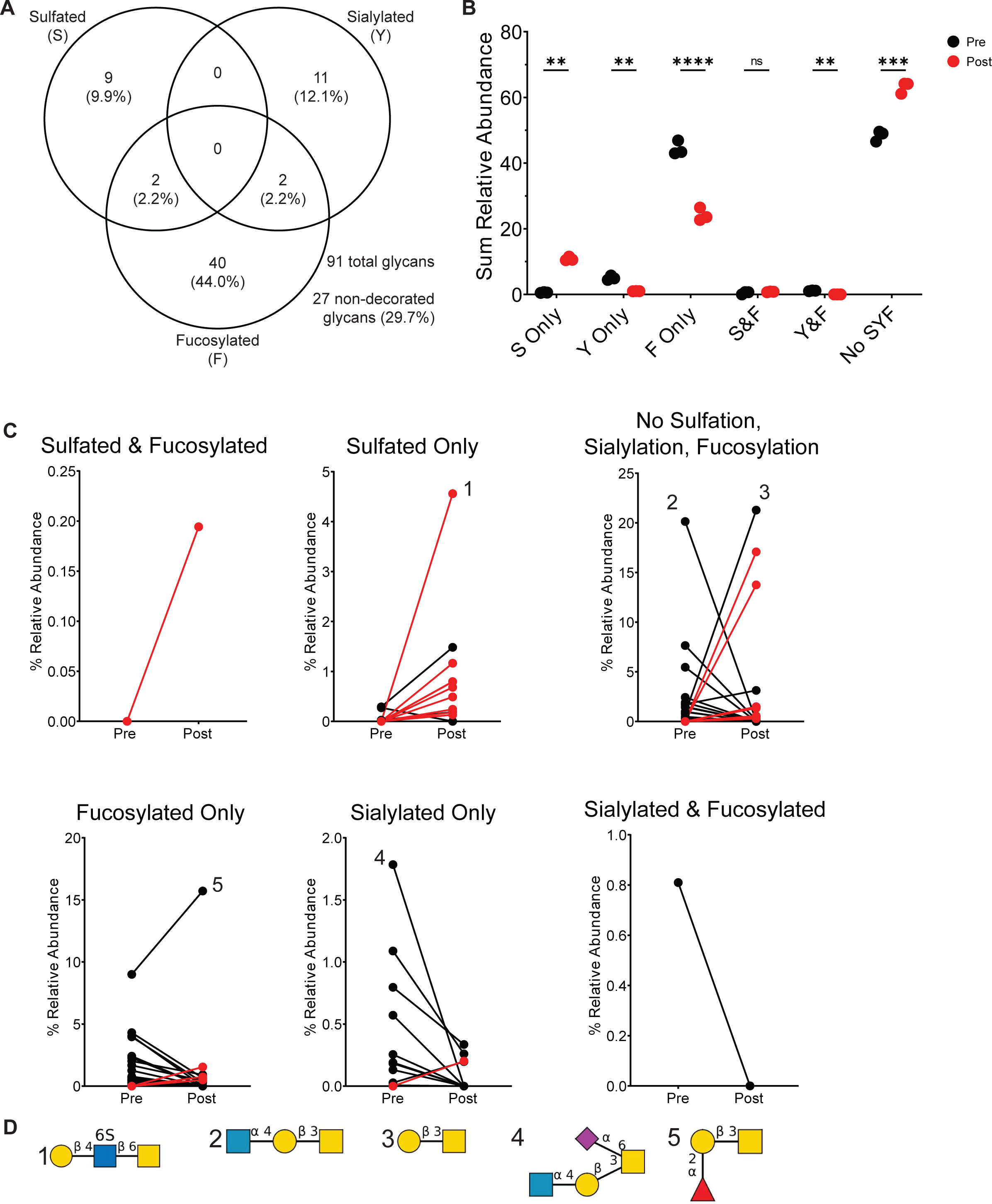
Fucosylated and sialylated glycans are degraded after growth of *R. torques* on PGM, but sulfated glycans accumulate. **A**, Number and percentage of total glycans detected in PGM pre- and/or post-growth of *R. torques* displaying indicated structural features by LC-MS/MS. **B**, Sum relative abundance of glycans and peeling reaction products pre- or post-*R. torques* growth on PGM with respective structural features. Statistical analysis was performed with paired, two-tailed t tests between pre- and post-growth samples for each structural feature category. S=sulfated, Y=sialylated, F=fucosylated. **p<0.01, ***p<0.001. **C**, relative abundance of glycans detected that were retained on the mucin polypeptide backbone and were significantly different in pre- or post-growth samples of *R. torques* on PGM. Red lines indicate glycans that were not detected in any of the pre-growth samples but were detected in the post-growth samples. Each point represents the average relative abundance of three samples. **D**, Putative structures of select detected glycans. Numbers refer to corresponding plot in C. Structures generated with DrawGlycan^63^.

**Supplemental Figure S11.**
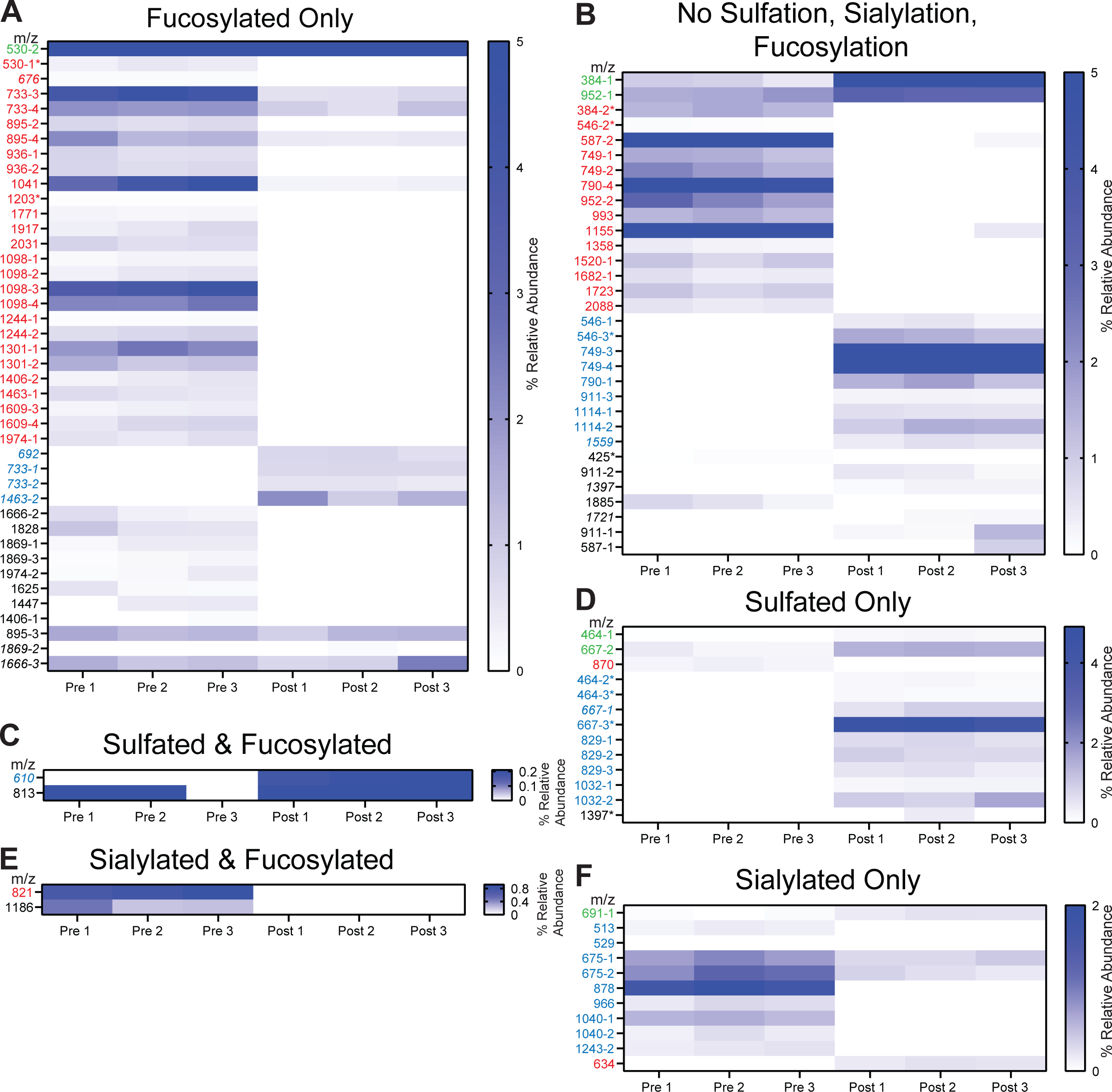
LC-MS/MS glycans identified before and after growth of *R. torques* on PGM. **A-F**, Glycans detected were grouped based on structural features: fucosylated only (**A**), no sulfation, sialylation, or fucosylation (**B**), sulfated and fucosylated (**C**), sulfated only (**D**), sialylated and fucosylated (**E**), and sialylated only (**F**). Heat maps indicate percent relative abundance of each glycan pre- or post-growth of *R. torques.* Molecular weights in green represent glycans detected in at least one pre-growth sample that significantly increase in abundance post-growth; glycans in red decreased in abundance post-growth compared to pre- growth, glycans in blue were undetected pre-growth but were detected post-growth; and glycans in black were not significantly different in abundance pre- or post-growth. Statistics were analyzed using paired, two-tailed t-tests between the three pre- and three post-growth samples for each glycan. MW = molecular weight. * = peeling reaction product, italicized m/z values = *N*- glycan.

**Supplemental Figure S12.**
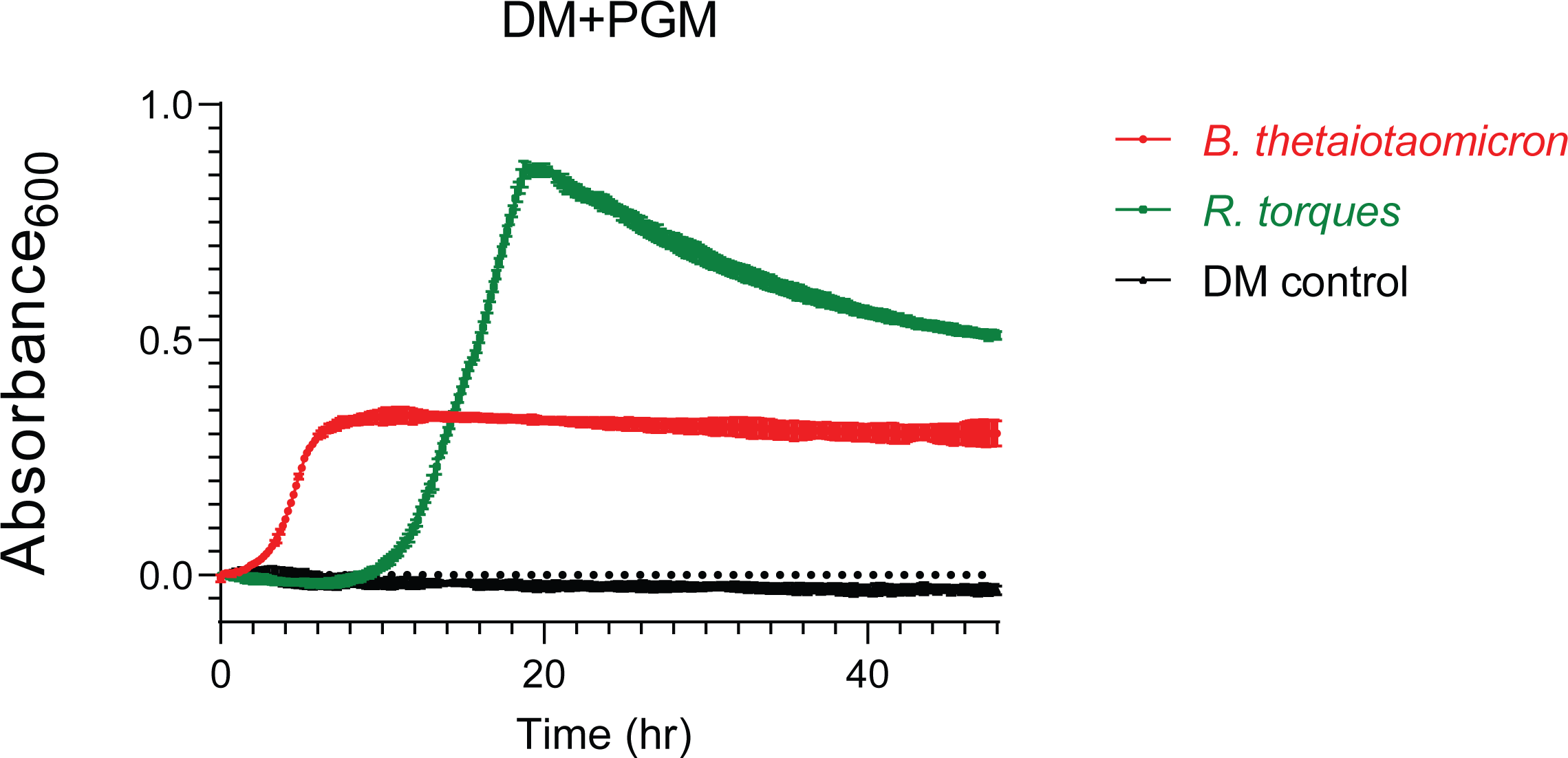
Growth of *B. thetaiotaomicron* and *R. torques* on PGM in partially defined medium (DM). *B. thetaiotaomicron* tdk^-/-^ and *R. torques* were grown anaerobically at 37°C in DM with PGM (10 mg/ml final concentration) as the major carbohydrate source and growth was monitored by Absorbance_600_. DM control contained DM+PGM only. Each point represents n=3 and error bars represent SD.

**Supplemental Figure S13.**
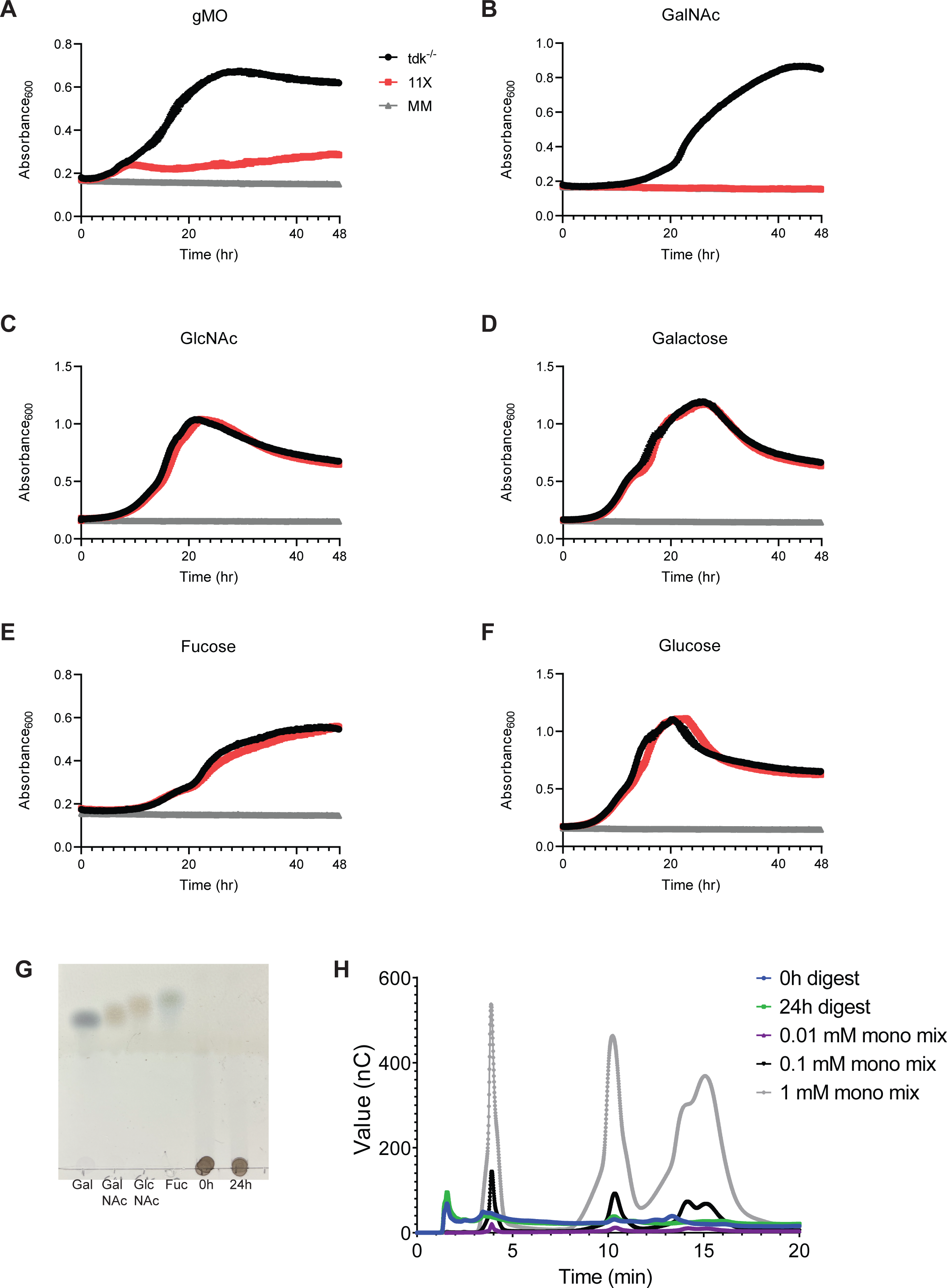
Growth of *B. thetaiotaomicron* tdk^-/-^ and *B. thetaiotaomicron* 11X mutant on glucose, MOG, and mucin monosaccharides and dialysis validation of *R. torques* PGM digest. **A-F**, Both strains of *B. thetaiotaomicron* were grown in Bacteroides minimal medium with gMO (**A**), *N*-acetylgalactosamine (**B**, GalNAc), *N*-acetylglucosamine (**C**, GlcNAc), galactose (**D**), fucose (**E**), or glucose (**F**). MM = Bacteroides minimal medium negative control. gMOs were at a final concentration of 10 mg/ml, and all others were 5 mg/ml. The first collected A_600_ value (0 h) was omitted from all conditions due to high baseline noise. Each point represents n=3 and error bars represent SD. **G**, thin layer chromatogram of *R. torques*-digested PGM substrates (0h control and 24h digest) to assess efficiency of dialysis in removing monosaccharides from these substrates. Standards at 10 mM were spotted once (3 µl), PGM substrates at 20 mg/ml were spotted twice (3 µl each). Gal=galactose, GalNAc=*N*-acetylgalactosamine, GlcNAc=*N*-acetylglucosamine, Fuc=fucose. **H**, HPAEC-PAD analysis of 0h and 24h PGM digests for presence of monosaccharides after dialysis. Mono mix=fucose (fuc), galactose (gal), *N*-acetylgalactosamine (GalNAc), and *N*-acetylglucosamine (GlcNAc). Concentrations for mono mixes reflect concentration of each monosaccharide in the mix.

**Supplemental Table 1. RNA-sequencing MOG vs. glucose fold change data, functional annotations of differentially expressed genes, RPKM in glucose and MOG.**

**Supplemental Table 2. Relative abundance of highly expressed proteins in the *R. torques* supernatant detected by LC-MS/MS across multiple proteomics runs and sample preparation types.**

**Supplemental Table 3. Top hits from proteomics and transcriptomics datasets from Venn diagram.**

**Supplemental Table 4. Composition and putative structures (if available) of glycans detected by LC-MS/MS analysis of samples pre- and/or post-growth of *R. torques* on cMUC2.**

**Supplemental Table 5. Composition and putative structures (if available) of glycans detected by LC-MS/MS analysis of samples pre- and/or post-growth of *R. torques* on PGM.**

